# Central Presynapses Regulate Spontaneous Synaptic Vesicle Exocytosis Rate by Constraining Recycling Pool Density

**DOI:** 10.1101/2024.09.19.613487

**Authors:** P. Wilson, N. Kim, R. Cotter, M. Parkes, M.N. Reed, M.W. Gramlich

## Abstract

Synapses represent a fundamental unit of information transfer during cognition. They accomplish this via presynaptic vesicle exocytosis, which can occur either spontaneously or by an action potential leading to evoked release. It has been well established that evoked release is probabilistic in nature, but it has been less clear what mechanisms mediate spontaneous release. Understanding spontaneous release is important because it is an essential maintenance mechanism for synaptic connections. We propose a mechanistic framework and model of spontaneous release based on immobile vesicles in the reserve pool geometrically constraining mobile vesicles in the recycling pool, which provides a force leading to a spontaneous release rate. We experimentally support this framework using a combination of Scanning Electron Microscopy (SEM), high-resolution fluorescence microscopy techniques using pHluorin-VGlut1 and a single vesicle SGC5 reporter, and a computational model. We observe that the spontaneous release rate increases linearly with the number of vesicles but is constant in the absence of presynaptic actin. We then use an acute agent, Forskolin, to further constrain the volume of the recycling pool, leading to an increased spontaneous release rate. We show that our framework predicts the increasing spontaneous release rate experimentally observed. These results suggest that synapses constrain the density of the recycling pool to mediate spontaneous release rate via the entropic force.

## Introduction

Synaptic transmission represents the basic unit of information transfer during cognition. This occurs predominantly between chemical synapses where neurotransmitter (NT) carrying synaptic vesicles (SVs) fuse with the plasma membrane (exocytosis) at the active zone (AZ) of the presynapse to release NT into the synaptic cleft.^1–5^ This process is efficiently maintained by a complex recycling process that keeps a large collection of SVs maintained in different pools (Reserve, Recycling, Ready releasable) within each presynapse.^6–14^ This complex process allows for presynapses to maintain mechanisms that provide a robust NT release in response to action potentials (APs) across a large range of stimulation conditions, from single release events ^2,4,15–17^ to high-frequency stimulation responses.^15,18–26^

In parallel with AP-induced evoked SV exocytosis, SVs will spontaneously exocytose in the absence of APs (spontaneous release).^13,16,17,27–29^ This spontaneous release has long been considered as a support process of multiple mechanisms for synaptic connections.^13,27,29^ Changes in spontaneous release have also corresponded to changes in synaptic transmission during learning and memory.^30–32^ Dysfunction in spontaneous release has been shown and proposed in multiple studies to correlate with neuronal diseases.^33–35^ These diverse range of studies have shown that spontaneous release is an essential process in the fundamental operations of synaptic transmission.

However, the mechanisms that regulate spontaneous release have been less clear as they do not correspond with the same mechanisms that regulate evoked SV exocytosis.^15–17,28^ Evoked SV exocytosis follows a well-established multinomial release model across a wide range of synapse types.^36–38^ This process involves a pool of SVs (RRP) tethered at individual release sites in the AZ ready to engage in AP-mediated exocytosis, followed by a larger pool of SVs (Recycling) that replace the exocytosed RRP SVs, and finally a large contained pool of SVs (Reserve) thought to maintain the Recycling pool. In this model framework, any given release site has a finite probability (P_r_) of releasing a single tethered RRP SV in response to an AP, and the total synaptic transmission response of the synapse is the number of release sites (n) times their individual release probability. Many different theoretical and experimental approaches have been successfully employed to support the multinomial model based on parameters such as the kinetic rates of different constituents (Ca^2+^, or the exchange of SVs between pools, etc.).^13–15,23,28,29,39,40^ This approach has been less successful in modeling and understanding the mechanics of spontaneous SV release. Studies have shown that some parameters correlate with spontaneous release, such as the size of the RRP,^16^ or internal constituent concentrations ^33^ but no single study has developed a single coherent framework to model and predict spontaneous release.

In this study, we develop a useful framework and model of spontaneous release as a holistic outcome of the total synaptic structure and organization of SV pools. In this framework, the reserve SV pool exists to regulate the density of the recycling pool. The SVs in the recycling pool then exert an entropic force at the SVs in the RRP. Spontaneous release rate is then proportional to the entropic force on the RRP. We introduce this framework using experimentally measured Large Area Scanning Electron Microscopy (La-SEM) of in vitro cultured primary hippocampal neurons combined with a computational model that relates the presynaptic structure and distribution of SV pools to spontaneous release rate. We then support this framework by directly measuring the spontaneous release rate and correlating it with the total SV pool size using the fluorescent reporter pHluorin-VGlut1. We show how the loss of the reserve pool, via loss of polymerized actin using Latrunculin-A, results in a loss of spontaneous release rate regulation. We then use SGC5 labeled single SV mobility within the presynapse combined with a computational model to show how recycling pool SV mobility leads to the spontaneous release rate observed with pHluorin-VGlut1. Finally, we use the agent Forskolin, which mimics memory formation and has been shown to induce a volume restriction on the Recycling pool,^41–44^ and we show leads to a further increase in spontaneous release rate. These combined experimental and computational results support the model that the presynaptic spontaneous release rate is regulated by the structure and distribution of SV pools. This entropic force model provides a coherent way to understand how complex changes occur during synaptic plasticity, memory formation, and neurodegeneration.

## Results

### Entropic Force Based Model of Spontaneous Release

To understand how vesicle pool size and distribution within the presynapse influence spontaneous synaptic transmission, we developed a framework of the spontaneous force exerted by SVs on exocytosis events at the active zone based on experimental SEM results. We cultured presynaptic hippocampal neurons with astrocytes for 14-21 DIV following our previously established approach (See Methods).^45,46,11,47^ We then fixed, stained and imaged cultures under homeostatic conditions using our previously established large-area scanning electron microscopy (La-SEM) approach (See Methods).^39^ This allowed us to distinguish a large distribution of individual presynapses with different sizes and numbers of SVs.

We quantified the relationship between presynaptic geometric size and number of SVs based on two different frameworks. First, we analyzed the total number of SVs in presynapses based on the perspective that every SV can freely traverse the entire presynapse volume (Unconstrained, Fig. 1A); in this framework, all SVs (which we will call the ***Total Pool***) contribute to the internal entropic force. Second, we analyzed the distribution of SVs as a function of distance from the presynaptic active zone, based on the now established internal geometric structure of presynapses (Constrained, Fig. 1B);^48^ in this framework, only the SVs closest to the presynaptic active zone (which is commonly called the ***Recycling Pool***) contribute to the entropic force (Green lines, **Fig. 1B**), while SVs farther from the active zone are constrained to each other and the actin cytoskeleton (which is commonly called the ***Reserve Pool***) preventing them from contributing to the entropic force (Red lines, **Fig. 1B**). Importantly, the contribution of entropic force to spontaneous release is also mediated by the size of the active zone (equivalent to the number of SVs at the active zone, which we will call ***RRP***) under both frameworks (Purple line, **Fig. 1B**).

**Figure 1:**
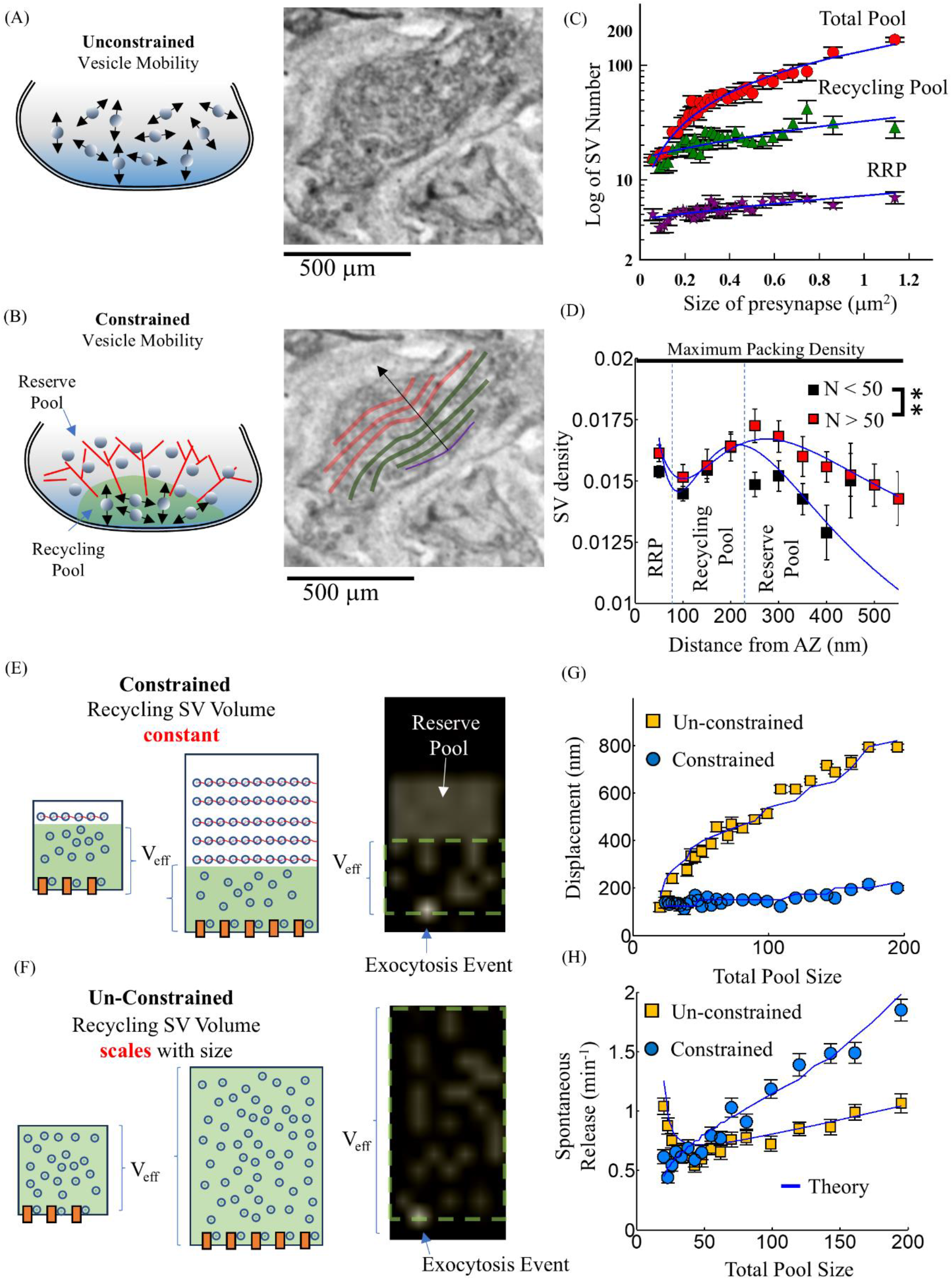
Entropic Force Framework Model Based on Experimental La-SEM Measurements (A) The unconstrained model of SVs traveling the entire size of the presynapse (Left panel) based on a simple analysis of counting the number of SVs in a presynapse using La-SEM (Right panel). (B) The constrained model of SVs assumes only the recycling pool is mobile (Left panel) and is based on a distance-based analysis of SVs relative to the presynaptic active zone (Purple line, Right panel). SVs in the recycling pool occur within the first 200 nm or less (Green Lines, Right panel). SVs in the reserve pool occur at distances greater than 200 nm (Red Lines, Right panel). (C) The number of SVs and their distribution within the presynapse increase linearly with presynapse size (data compared on log scale to show scale differences). The total number of SVs increases linearly (129 SVs per micron^2^) with the size of the presynapse (Blue Line). The number of SVs in the Recycling Pool (Green Triangles) also increases linearly but at a slower rate (17 SVs per micron^2^) with presynapse size (Blue Line). The SVs in the RRP closest to the active zone (Purple stars) increase very little (3 SVs per micron^2^) with presynapse size (Blue Line). (D) All presynapses exhibit the same overall SV density distribution which is high at the AZ (RRP), lower for the next 100 nm (Recycling pool) followed by a slightly higher density around 250 nm from the AZ (Reserve Pool). There is a slight increase in the density of SVs in the RRP and Recycling Pool for presynapses with greater than 50 SVs (Red Squares) compared to less than 50 SVs (Black Squares). There is a significant increase in Reserve Pool density for presynapses with greater than 50 SVs (Red Squares) compared to less than 50 SVs (Black Squares). (E) The constrained lattice based model (Top Panel) fixes the location of the Reserve Pool SVs starting at 250 nm from the active zone for all presynapse sizes. Only the Recycling pool SVs are allowed to move freely within the reduced effective volume (Green region). The un-constrained lattice based model (Bottom Panel) allows all SVs to move freely for all presynapse sizes so that the effective volume scales with presynapse size (Green region). (F) The constrained simulation model (Top panel) shows mobile SVs moving within a limited region and an exocytosis event is counted whenever a SVs reaches the active zone (z = 0) with a finite probability p_ψ_. The unconstrained simulation model (Bottom panel) shows mobile SVs moving within the entire presynapse and an exocytosis event is counted whenever any SVs reaches with active zone site with the same finite probability as the constrained model. (G) Total mean-square SV displacement for the un-constrained model (Yellow Squares) reaches a maximum roughly equal to the size of the presynapse (Blue line) during simulation. The mean-square SV displacement for the constrained model (Blue Circles) scales roughly with the size of the constrained volume of the recycling pool (Blue line). (H) The spontaneous release rate is calculated as the average number of registered exocytosis events per minute across all simulations for a given total SV number. The spontaneous rate increases linearly constrained model (Blue Circles) and follows the theoretical entropic force framework (Blue line). The spontaneous rate is relatively constant in the un- constrained model (Yellow Squares) and follows the theoretical entropic force framework (Blue line) assuming the volume scales linearly with presynapse size. N_presynapses_ = 98; N_simulations_ = 100 per SV number Data points represent average values from binned raw data with bins set to have fixed number of data points per bin. Statistics pair-wise 2-sampled t-Test.* = P< 0.05; ** = P < 0.01; *** = P< 0.001

These two different analysis approaches easily distinguish the relative difference in the number of SVs in each pool as a function of presynapse size. The total number of SVs (Red circles, **Fig. 1C**) follows a linear relationship with the size of the presynapse (Blue line, **Fig. 1C**). This result would suggest that the number of SVs almost doubles when the geometric size of the presynapse doubles (a relative increase of 1.8 in SV number when the presynapse size doubles). However, the number of the SVs in the recycling pool (Green triangles, **Fig. 1C**) increase at a significantly slower linear rate (Blue line, **Fig. 1C**). Last, the number of SVs at the active zone (Purple stars, **Fig. 1C**) remains relatively constant with presynapse size reaching a maximum when the total number of SVs is greater than 50. These results are consistent with the well-established understanding of SV pool distributions; however, these results do not distinguish if or how the density of the SV pools changes with presynapse size, which would have significant effects on entropic force.

To determine if the density of SVs changes with presynapse size, we quantified the density of SVs relative to the active zone independent of any pool distribution. We quantified the radial density of SVs as a function of distance from the AZ (Black arrow, **Fig. 1B**, See Methods) in increments of 50 nm, which is roughly the diameter of an SV.^49^ We observed that regardless of size, SV density followed the same general relationship: it is high at the active zone (x = 50 nm, **Fig. 1D**); then decreases for the next 100 nm from the active zone; followed by increasing density to a maximum at approximately 200-250 nm from the AZ (**Fig. 1D**); and, SV density decreases again starting at 250 nm from the active zone.

The overall scale of the distribution in SV density does change with presynapse size. Presynapses with less than 50 total SVs (Black squares, **Fig. 1D**) have less density overall than presynapses with greater than 50 SVs (Red squares, **Fig. 1D**). More importantly, as presynapse size increases, the majority of increasing SV density occurs at larger distances from the active zone (>250 nm, **Fig. 1D**). These results are consistent with the relative change in SV number results, as well as the now established geometric structure of the presynaptic cytoskeleton.^48^ These results suggest that *the structure of the presynapse uses the reserve pool itself to constrain the recycling pool density*, which would have significant effects on how the entropic force of the recycling pool would mediate the spontaneous release rate.

Using the experimentally observed distribution of SVs within the presynapse, we propose the following model based on an entropic force framework (See Methods for derivation):

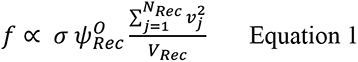

where the spontaneous release rate ***f*** is mediated by the volume of the recycling pool V_Rec_, the number of active zone release sites σ, the number of SVs in the recycling pool (N_rec_), proportional to their kinetic energies 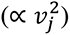, and a spontaneous correlation parameter 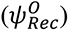.

The **spontaneous correlation parameter** 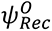 represents the native spontaneous release rate per occupied release site. This is fundamental to the structure, function, and local environment of release sites themselves.

This model predicts that as the number of SVs within the presynapse increases, the recycling pool SVs are increasingly constrained by the reserve pool, resulting in an increasing density of recycling SVs. The increased SV density, in turn, leads to increased entropic force at the AZ, resulting in an increasing spontaneous release rate.

In order to show how this constrained volume of the recycling pool within the presynapse leads to changes in spontaneous release, we developed a computational model of SV mobility within the presynapse for different possible geometries (See Methods for model derivation). We first modeled the constrained geometry where the reserve pool SVs act as a fixed barrier for the recycling pool SVs reducing their effective volume (Constrained, **Fig. 1E**). The number of SVs in the recycling pool and reserve pool follow the relationship observed in SEM results (**Fig. 1C**). Alternatively, we modeled an unconstrained geometry without a reserve pool where all SVs move freely (Unconstrained, **Fig. 1 F**). In this model, a spontaneous release event occurs with a finite probability (p_ψ_) whenever an SV hits an AZ release site. The rate of spontaneous release is then calculated as the average number of exocytosis events measured per minute. Finally, we model the number of AZ release sites as proportional to the volume corrected number of RRP SVs (**Fig. 1C**):

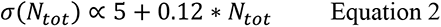

This computational model supports our hypothesized entropic force theoretical framework. First, the constrained model shows a relatively constant SV displacement regardless of SV pool size (Constrained, **Fig. 1G**), while the unconstrained model shows an increasing SV displacement proportional to the radius of the presynapse (Unconstrained, **Fig. 1G**). This differential motion is consistent with the hypothesis that the reserve pool constrains the recycling pool mobility. Importantly, the constrained model shows an increasing spontaneous release rate with increasing total SV pool size (Constrained, **Fig. 1H**); whereas the unconstrained model shows a relatively constant spontaneous release rate (Unconstrained, **Fig. 1G**). Further, the theoretical entropic force framework predicts both the constrained and unconstrained computational model results (Solid blue lines, **Fig. 1G**).

These combined La-SEM, theoretical, and computational results show that the presynaptic spontaneous release rate can be mediated by the density of the recycling pool. For the remainder of this study, we will experimentally test each predicted parameter (SV number, SV mobility, spontaneous release rate) using different approaches to support this framework.

### Spontaneous Release Rate Increases with Total Synaptic Vesicle Pool Size

To determine if the entropic force framework can predict experimental presynaptic spontaneous release, we quantified presynaptic release mechanics using a pH-sensitive fluorescent reporter attached to the vesicular glutamate reporter 1 (VGlut1). This reporter is dark inside the low pH environment of the vesicle but becomes fluorescent when the vesicle is exposed to the extracellular environment during exocytosis (**Fig. 2A**). This approach has a sensitivity range from single release events up to the total size of the synaptic vesicle pool. We, and others, have previously used the VGlut1-pHluorin reporter to distinguish spontaneous release mechanics,^17,50,51^ stimulated release mechanics,^19,45,50–52^ active zone size,^2^ recycling pool size,^45,51^ and total synaptic vesicle pool size.^45^

**Figure 2:**
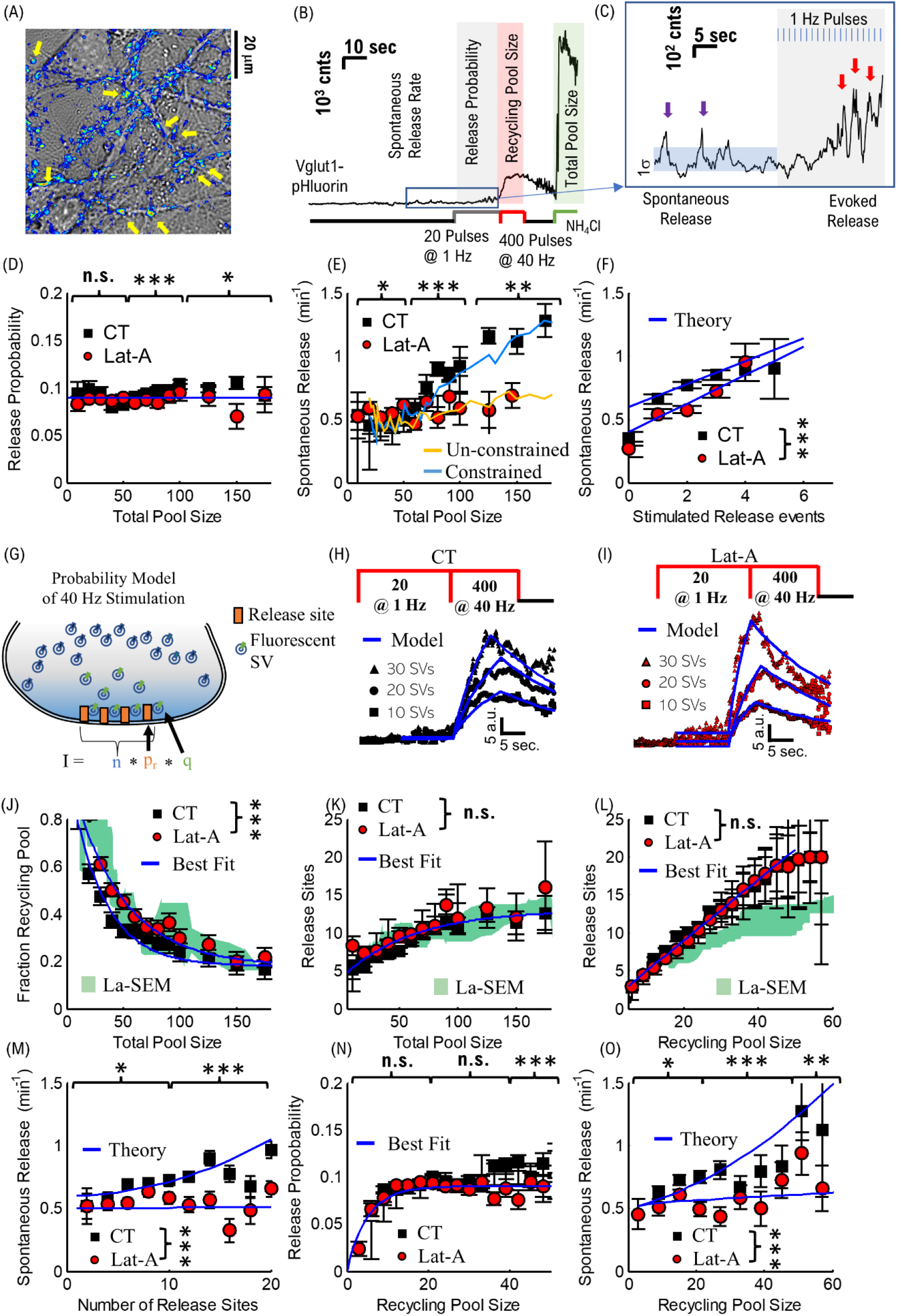
**Experimental Spontaneous Release Rate in pHluorin-VGlut1 labeled Hippocampal Presynapses Support Entropic Force Theory** (A) Transmission image (Grey scale) of cultured hippocampal neurons labeled with pHluorin-VGlut1 exhibit intensity at presynaptic locations (Blue scale) when stimulated. (B) An example of the stimulation protocol used that quantifies pHluorin-VGlut1 intensity (i) in the absence of stimulation to measure spontaneous release events, (ii) with 1Hz stimulation to measure evoked release probability, (iii) with 40Hz stimulation to measure total recycling pool size, (iv) under exposure to NH_4_Cl to measure total SV pool size. (C) An example of single SV release events counted for spontaneous release (Green arrows) and evoked release (Red arrows) events. Events are counted when the intensity increases at least 1 standard deviation above the noise within at least 0.5 sec time-window and for at least 1.0 sec. (D) The release probability as a function of total pool size. The average release probability is constant for all SV pool sizes (∼ 0.1) for both control (CT, Black Squares) and presynapses exposed to Latrunculin-A (Lat-A, Red Circles). (E) The spontaneous release rate measured as a function of total SV pool size increases linearly for control presynapses (Black Squares) but is constant for Lat-A presynapses (Red Circles). The constrained model (Blue line) reproduces the control results, and the un-constrained model (Yellow line) reproduces the Lat-A results. (F) The spontaneous release rate increases linearly as a function of evoked release events for both control (Black squares) and Lat-A (Red circles) presynapses. Control presynapses exhibit an overall larger spontaneous release rate than Lat-A presynapses. Both results follow the entropic force framework theory (Blue lines) assuming the number of stimulated release events correlate with the number of release sites and number of SVs in the recycling pool (panel L). (G) The 40Hz pHluorin-VGlut1 intensity is modeled using the established multinomial probability theory based on the number of release sites (**n**) the probability of release per site (**p_r_**) and the intensity per SV (**q**). The number of release sites is variable, while the release probability and intensity per SV are obtained from experimental measurements (D, and C respectively) (H) Example traces of control presynapse pHluorin-VGlut1 intensity during 40Hz stimulation fit using the multinomial model (Blue lines) for 10, 20, 30 SVs. (I) Example traces of Lat-A presynapse pHluorin-VGlut1 intensity during 40Hz stimulation fit using the multinomial model (Blue lines) for 10, 20, 30 SVs. (J) The fraction of SVs in the Recycling pool relative to the total number of SV pool size. Both control (Black Squares) and Lat-A (Red Circles) data exhibit an exponential decrease (Blue lines) in the fraction of Recycling SVs with total pool size. The pHluorin-VGlut1 results also matched the measured La-SEM fraction (Green shaded region). (K) The number of release sites as a function of total pool size determined from multinomial model fit results increases linearly with total pool size up to ∼80 SVs followed by a constant number (n ∼ 10) as a function of total pool size (Blue line). The number of release sites is consistent with the number of SVs in the RRP measured with La-SEM (Green shaded region). (L) The number of release sites as a function of Recycling pool size determined from multinomial model fit results increases linearly with number of SVs (Blue line). The linear increase is ∼0.7 release sites per SV in the Recycling pool. (M) The spontaneous release rate increases linearly with the number of release sites for control presynapses (Black Squares) but remains constant in the presence of Lat-A (Red Circles). The entropic force theory (Blue lines) reproduces spontaneous release rates taking into account the average recycling pool size (not shown). (N) Release probability increases with increasing recycling pool size for up to 10 SVs for both control presynapses (Black Squares) and in the presence of Lat-A (Red Circles). The release probability then remains constant (0.1) for recycling pool size greater than 10 SVs. (O) The spontaneous release rate increases with recycling pool size for control presynapses (Black Squares) but remains constant in the presence of Lat-A (Red Circles). The entropic force theory reproduces both conditions when taking into account the average density of the recycling pool (not shown). CT: N_presynapses_ = 2057, from 6 different cultures; N_simulations_ = 100 per SV number Lat-A: N_presynapses_ = 1705, from 6 different cultures; N_simulations_ = 100 per SV number Statistics pair-wise 2-sampled t-Test.* = P< 0.05; ** = P < 0.01; *** = P< 0.001. Data points represent average values from binned raw data. Bins were set at fixed values to obtain a minimum of 30 data points per bin. Error bars represent SEM of binned data.

Here we developed a single imaging protocol that combines all previous measurements from the single release events to total pool size, but on a synapse-by-synapse basis (**Fig. 2B**). First, we image neurons in the absence of stimulation to capture spontaneous release frequency (**Fig. 2C**). Second, we electrically stimulate neurons at 1 Hz to measure evoked release probability (**Fig. 2C**). Then we stimulate the same neurons with 400 pulses at 40 Hz to measure the recycling pool (**Fig. 2B**). Finally, we expose neurons to NH_4_Cl to measure the total number of vesicles within the presynapse. This combined approach allows us to directly measure and relate multiple parameters (spontaneous release, evoked release, recycling pool size, total pool size) in the same synapse to distinguish the contribution of the vesicle pool size on presynaptic release mechanics.

We first determined the total pool size in each presynapse as the total intensity measured during NH_4_Cl exposure (ΔF_NH4Cl_) divided by the average single vesicle release intensity measured during spontaneous release (ΔF_single_). Our observed average pool size (N ∼ 70) is consistent with previously measured pool sizes and consistent with the volume corrected pool sizes measured with La-SEM (Supplementary Fig. S1 A).^53^ Further, the distribution of pool sizes results show that both pHluorin-VGlut1 and La-SEM approaches provide consistent and overlapping measures of total pool size.

To determine if the entropic force framework models presynaptic release, we compared experimentally measured spontaneous and evoked release mechanics as a function of the total synaptic vesicle pool size. We observed that evoked release probability is on average p_r_ ∼ 0.9 +/- 0.002 and independent of total pool size (Black squares, **Fig. 2D**) and consistent with previously measured release probability (p_r_ between 0.07 to 0.1).^19,45^ Alternatively, we observed that the spontaneous release frequency increases linearly with increasing total pool size (Black squares, **Fig. 2E**). This linearly increasing spontaneous release matches the theoretical entropic force framework theory and computational model (solid blue line, **Fig. 2E**). These results support our hypothesis that the volume of the recycling pool is constrained leading to an increasing spontaneous release frequency via increased SV density.

To test whether the reserve pool alters the effective volume and/or contributes to the spontaneous release directly, we acutely altered the actin network and measured presynaptic release mechanics. We exposed cells to the agent Latrunculin-A (Lat-A), which has been well established to drive actin toward depolymerization, resulting in a lower presynaptic vesicle reserve pool but still maintaining vesicles in the presynapse.^54^ We observed a slight reduction (∼6%) in evoked release for Lat-A (p_r_ = 0.088 +/- 0.002, Red Circles, **Fig. 2D**) compared to control (CT: p_r_ = 0.09 +/- 0.002). However, this limited change is consistent with the interpretation that loss of actin does not alter the recycling pool or mechanics of release sites at the active zone, as previously observed,^55–57^ and more likely increases the mobility of the Recycling Pool away from the active zone. We did, however, observe a significant change in spontaneous release frequency in the presence of Lat-A (Red Circles, **Fig. 2E**), compared to the control. Spontaneous release frequency became independent of the total vesicle pool size and remained constant at ∼0.5 SV/min. The constant spontaneous release rate also matches the unconstrained computational model of a constant SV number with volume (solid yellow line, **Fig. 2E**) based on our theoretical framework. These results support the hypothesis that the actin network serves to tether vesicles in the reserve pool and reduce the effective volume within the presynapse.

We finally compared the spontaneous release rate to the number of stimulated release events to determine if release probability correlated with spontaneous release. We observed the same linear increase in spontaneous release rate with increasing stimulated release for both CT (Black squares, **Fig. 2F**) and Lat- A (Red Circles, **Fig. 2F**) up to 4 released SVs (P_r_ ∼ 0.2), but the Lat-A rate was overall lower than for CT. We compared the measured spontaneous release rate to the theoretical rate (Eqn. 1, and Blue lines, **Fig. 2F**) estimated based on the average number of SVs (**Supplementary Figure S1 B**) and the number of release events multiplied by 5 as equivalent to the number of release sites. These results suggest that release probability at the active zone is a mediating factor in spontaneous release rate, assuming all other factors are constant, which is predicted in our entropic force framework via increasing release site number (σ).

These combined results support the hypothesis that the reserve pool functions to constrain the volume of the recycling pool to mediate spontaneous release frequency

### Release Site Number and Total Recycling Pool Size Are Secondary Mediators of Spontaneous Release Rate

It has been established that active zone size affects evoked release,^19,36^ and correlates with spontaneous release,^16^ and our observed correlation between spontaneous release rate and stimulated release events (**Fig. 2F**) supports this relationship. However, it is not known how the active zone size and recycling pool size combine to result in our experimentally observed spontaneous release rate (**Fig. 2E**). To determine how the recycling pool dynamics and/or presynaptic release mechanics affect spontaneous release rates, we modeled VGlut1-pHluorin intensity during 40 Hz stimulation (**Fig. 2G**). We used the established multinomial release probability model,^36,37,58^ which we have previously used to extract the number of vesicles in the recycling pool as well as the number of release sites (See methods).^45^ Briefly, we used experimental values (intensity per vesicle and evoked release probability) as fixed parameters in the simulation. We then fit the experimental 40 Hz pHluorin-VGlut1 intensity (**Fig. 2H**) to simulated intensities (Solid Blue lines, **Fig. 2H**) of different parameters (number of release sites, endocytosis rate, start/end release times) to determine the number of vesicles released during stimulation as well as the number of release sites. This approach allowed us to combine the number of recycling pool SVs, number of release sites, and release mechanics in the same synapses that we determined spontaneous/evoked release events as well as total vesicle pool size (**Fig. 2B**).

Using this model approach, we observed that the fraction of the recycling pool decreased exponentially with the total number of vesicles (Black Squares, **Fig. 2J**), consistent with the same relationship observed using La-SEM (Shaded Green region, **Fig. 2J**). When compared to the total pool size, this result suggests that the recycling pool is predominantly around 20 vesicles or less. Further, this result supports the hypothesis that the majority of vesicles in the synapse are tethered in the reserve pool while the recycling pool size is maintained at a constant size for the majority of presynapses (N_tot_ < 70 SVs). Importantly, we observed that the relationship between the recycling pool size and total number of vesicles was not significantly changed in the presence of Lat-A (Red circles, **Fig. 2J**). This suggests that the number of vesicles engaging in synaptic transmission is not regulated by the total number of vesicles within the presynapse.

We then compared the number of release sites to the total pool size (**Fig. 2K**) to determine if active zone size correlates with total pool size in the same presynapses as increasing spontaneous release rates. We modeled each release site as having the same mean release probability mechanics (p = 0.01) based on experimentally observed release probabilities (**Fig. 2D**) and allowed only the number of release sites to change (See Methods). Initially, the number of release sites increases linearly with the total pool size for both CT and Lat-A, but then the average number of release sites remains constant for total pool sizes greater than 100. This is consistent with La-SEM measurements of the number of SVs observed in the RRP (**Fig. 1C**) after volume corrected (Shaded Green region, **Fig. 2K**).

We then combine the release site results into the entropic force framework as:

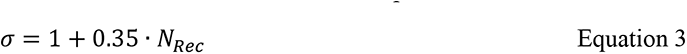

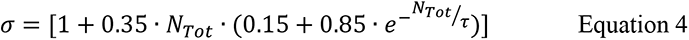

We also compared the number of release sites as a function of the recycling pool size (**Fig. 2L**) because it is well established that SVs in the RRP dynamically exchange with SVs in the Recycling pool and thus not every release site is occupied at all times.^4,5,16,59^ We observe a linear increase (Blue Line, **Fig. 2L**) in the number of release sites with the total number of SVs in the recycling pool for both control (Black Squares, **Fig. 2L**) and Lat-A (Red Circles, **Fig. 2L**). The number of release sites increases at roughly half the rate as the total number of SVs in the recycling pool (i.e. a release site gets added for every two additional SVs in the Recycling pool). However, the pHluorin-VGlut1 results show a larger number of release sites than the number of RRP SVs measured using La-SEM (Shaded Green region, **Fig. 2L**). These differences are consistent with the established observation that not every release site is occupied which would result in a lower number of measured RRP SVs (measured with La-SEM) than the number of available release sites (measured by pHluorin-VGlut1).

To determine if the number of release sites correlates with spontaneous release rate, we compared the experimentally measured spontaneous release rate to the fit number of release sites from the 40 Hz model (Black squares, **Fig. 2M**). Spontaneous release rate increases continuously with release sites up to 7-10 sites, consistent with recently observed correlations,^16^ but does not increase at the same rate as observed for the total pool size (**Fig. 2E**). One important reason that spontaneous release rate is less correlated with release site number is that a set number of release sites does not mean a set number of Recycling Pool SVs; Indeed, we used the entropic force model (**eqn. 1**), but converted the number of release sites to the equivalent total number of SVs (**Supplementary Fig. S1 C, D**), and found the number of release sites still correlated with the equivalent spontaneous release rate (Blue Line, **Fig. 2M**).

We then compared the spontaneous release rate to release sites in the presence of Lat-A (Red circles, **Fig. 2M**) to support our hypothesis that the reserve pool is the mediating factor. We observed the spontaneous release rate is relatively flat and independent of the number of release sites, suggesting that the increase observed for the CT data is due to the reserve pool constraining the volume of the recycling pool rather than the number of release sites directly contributing.

We next compared evoked release probability to the number of vesicles in the recycling pool to determine if the size of the recycling pool correlated with vesicle release. We observed that evoked release probability increases with recycling pool size up to 15-20 vesicles (Black squares, **Fig. 2N**), but then remains constant above 20 vesicles. This relationship was the same in the presence of Lat-A (Red Circles, **Fig. 2N**). These results suggest that loss of actin does not affect vesicle release mechanics at the active zone, as consistent with previous findings.^56^ Further, the change in spontaneous release frequency observed in the presence of Lat-A (Red Circles, **Fig. 2 F,M**) is not due to changes in release mechanics at the active zone. This result is also consistent with previous studies showing that evoked release and spontaneous release are unrelated phenomena.^15^

Last, we compared the spontaneous release frequency with the recycling pool size to determine if the recycling pool size alone mediated observed spontaneous release rates. We observed that the spontaneous release rate shows an increase with recycling pool size (Black squares, **Fig. 2O**). This rate was slower than for the spontaneous release rate as a function of total SV pool size (**Fig. 2E**). This difference is due to the fact that a given recycling pool size does not mean a set SV density. Indeed, when estimating the fraction of Recycling SVs and equivalent total pool size (**Supplementary Fig. S1 E,F**), the resulting theoretical spontaneous release rate matches the experimentally observed result (Blue line, **Fig. 2O**). Alternatively, we observed a constant spontaneous release frequency in the presence of Lat-A up to 40 vesicles (Red circles, **Fig. 2O**). These results suggest that the recycling pool size alone does not significantly alter spontaneous release rates. Rather, these results support the hypothesis that the constrained volume of the recycling pool mediates spontaneous release frequency.

Taken together, these results support the framework that the observed spontaneous release rate is predominantly mediated by the constrained density of the recycling pool SVs and not directly mediated by the active zone mechanics itself.

### Synaptic Vesicle Mobility Supports Entropic Force Framework

We sought to support the pHluorin-VGlut1 results with a second approach that could also test if vesicle mobility distributions supported the entropic force framework. To do this, we used an established fluorescence microscopy approach to label vesicles with a lipophilic reporter SGC5.^60^ We have previously shown that this approach can be used for single vesicle labeling to quantitatively distinguish recycling pool dynamics (**Fig. 3A-C**).^39^ Further, using this approach in combination with a correlation analysis algorithm (See methods),^11,46,47^ we can quantify vesicle speeds and positions within presynapses to distinguish molecular level changes (**Fig. 3D,E**).

**Figure 3:**
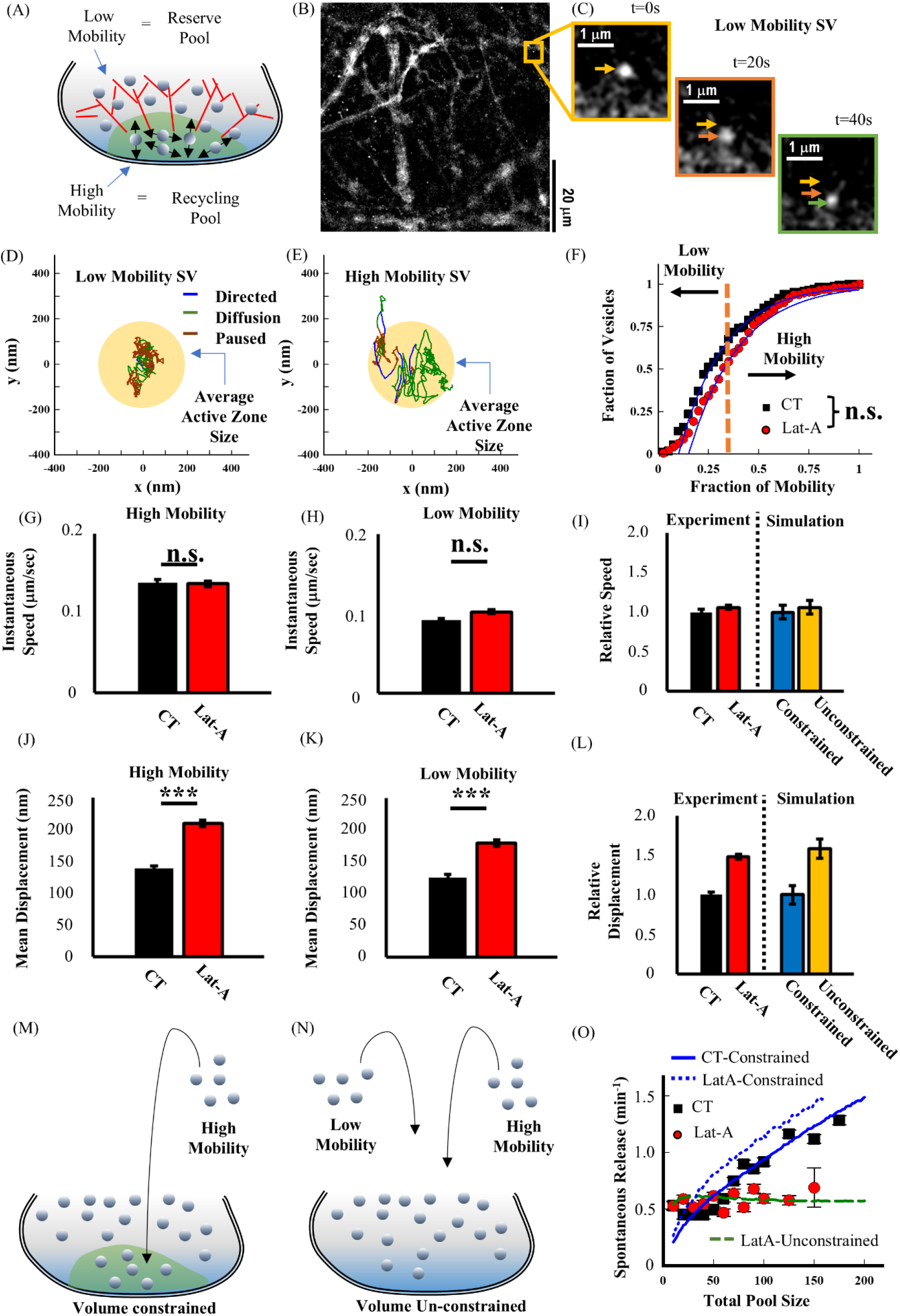
Experimental Single SGC5 Labeled Synaptic Vesicles Reproduce Spontaneous Release Rates and Support Entropic Force Theory (A) Cartoon Model of SV mobility distribution within presynapses. Low mobility SVs are presumed to be in the Reserve pool, and high mobility SVs are presumed to be in the recycling pool. (B) Example of Hippocampal Cell Cultures with SGC5 labeled single SVs. (C) Example of single SGC5 labeled SV mobility within a presynapse (D) Example of Low Mobility Single SV track showing bouts of Directed (Blue), Diffusive (Green), and Pausing (Red) behavior determined from correlation analysis algorithms. The overall mobility is compared to the average active zone size (Yellow Circle). (E) Example of High Mobility Single SV track exhibiting greater Directed and Diffusive motion compared to low mobility SVs (D) and compared to the average active zone size. (F) The distribution of SV mobility fraction measured as the total amount of time mobile (Directed + Diffusive) divided by the total amount of time observed. SVs in the presence of Lat-A (Red Circles) exhibit an increase in the fraction of time mobile compared to control SVs (Black Squares). (G) Average High Mobility SV speed is compared for control (CT, Black) and in the presence of Lat-A (Lat-A, Red). (H) Average Low Mobility SV speed is compared for control (CT, Black) and in the presence of Lat-A (Lat-A, Red). (I) Relative difference in speed for High Mobility SVs is compared for control (CT, Black) and in the presence of Lat- A (Lat-A, Red). The relative speed difference is also compared for the computational simulation models for constrained (Blue) and un-constrained (Yellow). (J) Average High Mobility SV rms-displacement is compared for control (CT, Black) and in the presence of Lat-A (Lat-A, Red). (K) Average Low Mobility SV rms-displacement is compared for control (CT, Black) and in the presence of Lat-A (Lat-A, Red). (L) Relative difference in displacement for High Mobility SVs is compared for control (CT, Black) and in the presence of Lat-A (Lat-A, Red). The relative displacement difference is also compared for the computational simulation models for constrained (Blue) and un-constrained (Yellow). (M) Cartoon representation of bootstrapping approach for the *volume constrained* model. SV tracks from the High Mobility group are randomly chosen and added to the presynapse in the Recycling pool. The number of SVs chosen is determined by the fraction of Recycling pool SVs measured with pHluorin-VGlut1 (Fig. 2J). (N) Cartoon representation of bootstrapping approach for the *volume constrained* model. SV tracks from the High and Low Mobility groups are randomly chosen and added to the presynapse in the total pool. (O) The virtual bootstrapped CT tracks in the volume constrained model (Black Solid Line) reproduces pHluorin- VGlut1 spontaneous release results (Black Squares). The virtual bootstrapped Lat-A tracks in the volume constrained model (Red Solid Line) over-estimates the pHluorin-VGlut1 spontaneous release results (Red Circles). However, the virtual bootstrapped Lat-A tracks in the volume unconstrained model (Red Dashed Line) reproduces the pHluorin- VGlut1. CT: N_SVs_ = 171, from 5 different cultures; N_simulations_ = 100 per SV number Lat-A: N_SVs_ = 193, from 4 cultures; N_simulations_ = 100 per SV number Statistics: 2-sampled KS-test performed in (F). For all other panels, pair-wise 2-sampled t-Test.* = P< 0.05; ** = P < 0.01; *** = P< 0.001. Data points represent average values separated SV (High/Low mobility) groups. Error bars represent SEM of separated data.

We first compared SVs based on their time spent mobile during observation (**Fig. 3F**) to determine changes in the distribution of SVs within the presynapse. We defined mobility as the fraction of time spent in directed (Blue, **Fig. 3D,E**) or diffusive (Green, **Fig. 3D,E**) motion as compared to pausing (Red, **Fig. 3D,E**). Most of the recently recycled SVs labeled with SGC5 exhibited low mobility (defined as less than 40% of time), and a minority of vesicles exhibited high mobility, defined as greater than 60% of time (Black, **Fig. 3F**). We observed the distribution of SV mobility increased slightly in the presence of Lat-A (Red, **Fig. 3F**). This increase in SV mobility in the presence of Lat-A supports the hypothesis that reserve pool SVs are no longer tethered to the actin network and allow recently recycled SVs more mobility within the presynapse.

We used the distribution of SV mobility to distinguish SVs as either in high or low mobility group. We then determined if SV dynamics in the high/low distributions changed in the presence of Lat-A. We observed that SV speed did not significantly change in the presence of Lat-A for the high mobility group (**Fig. 3G**) or the low mobility group (**Fig. 3H**). This result is consistent with the constant speed in the constrained versus unconstrained computational model (**Fig. 3I**). Alternatively, we observed that SV displacement significantly increased in the presence of Lat-A for both the high and low mobility group (Red, **Fig. 3J,K**) compared to the CT group (Black, **Fig. 3J,K**). These experimental results are consistent with the relative displacement changes in the unconstrained model relative to the constrained model (**Fig. 3L**).

These results support the hypothesis that loss of actin increases SV mobility within the presynapse by removing the reserve pool, thus increasing the available volume for recycled SVs to move.

### Bootstrapped Virtual Presynapses Using Vesicle Mobility Reproduces Spontaneous Release

The single SGC5 vesicle results do not distinguish if or how SV mobility and number combine to mediate spontaneous release rate observed in VGlut1-pHluorin results. To determine if single SGC5 results can reproduce spontaneous release rate results, we used a virtual bootstrap approach. This approach uses the entropic force framework to combine single vesicle SGC5 tracks to reproduce spontaneous release frequency (See methods). Briefly, we assumed tracks from the low mobility group represented SVs in the reserve pool (**Fig. 3A**), and high mobility tracks represent SVs in the recycling pool (**Fig. 3A**). We then randomly selected N-tracks (N = 10, 20, 30, etc…) from each distribution and extracted their average speeds. We then put the SV speeds into the entropic force equation (eqn. 1) and multiplied each by the **spontaneous correlation parameter** value. We assumed two possible geometries, bootstrapped SVs were either constrained in the volume via the reserved pool (**Fig. 3M**) or were unconstrained in volume equivalent to the presynapse size (**Fig. 3N**). Finally, we compared the inferred spontaneous release rates from the bootstrapping approach with the measured pHluorin-VGlut1 spontaneous release rate.

We compared the volume-constrained bootstrapped group of high-mobility SVs to the CT and Lat- A pHluorin-VGlut1 spontaneous release rate data (**Fig. 3O**). We observed that the CT bootstrapped data (Blue Solid Line, **Fig. 3O**) reproduced the pHluorin-VGlut1 spontaneous release rate (Black Squares, **Fig. 3O**) in the volume-constrained assumption. However, the Lat-A bootstrapped data (Red Solid Line, **Fig. 3O**) significantly overestimated the pHluorin-VGlut1 spontaneous release rate (Red circles, **Fig. 3O**) in the volume-constrained assumption. Alternatively, the same bootstrapped Lat-A data in the volume unconstrained assumption (Red Dashed Line, **Fig. 3O**) successfully reproduced the pHluorin-VGlut1 spontaneous release rate.

These bootstrapped results support the entropic force framework hypothesis that the spontaneous release rate increases with increasing SV pool size and dynamics under the condition that the recycling pool volume is constrained. Further, these results support the hypothesis that the actin network and reserve pool constrain the volume over which the recycling pool SVs traverse within the presynapse.

### Forskolin Increases Spontaneous Release Frequency via Reduced SV Recycling Pool Volume

In order to determine if the entropic force model can predict changes in spontaneous release rates from changes in SV pool dynamics, we exposed neurons to the acute pharmacological agent Forskolin. Forskolin has been established to mimic memory formation by inducing synaptic plasticity changes similar to those observed during long-term potentiation (LTP).^41–44^ Specifically, it has been shown that exposure to Forskolin causes dynamic changes in the recycling pool distribution causing SVs to remain closer to the active zone.^41^ Further, these changes in SV recycling pool dynamics correlate with changes in synaptic transmission.^30^ If Forskolin brings the SV recycling pool closer to the active zone, then our entropic force framework predicts that this should increase the density of the recycling pool (**Fig. 4A**), thereby increasing the spontaneous release rate.

**Figure 4:**
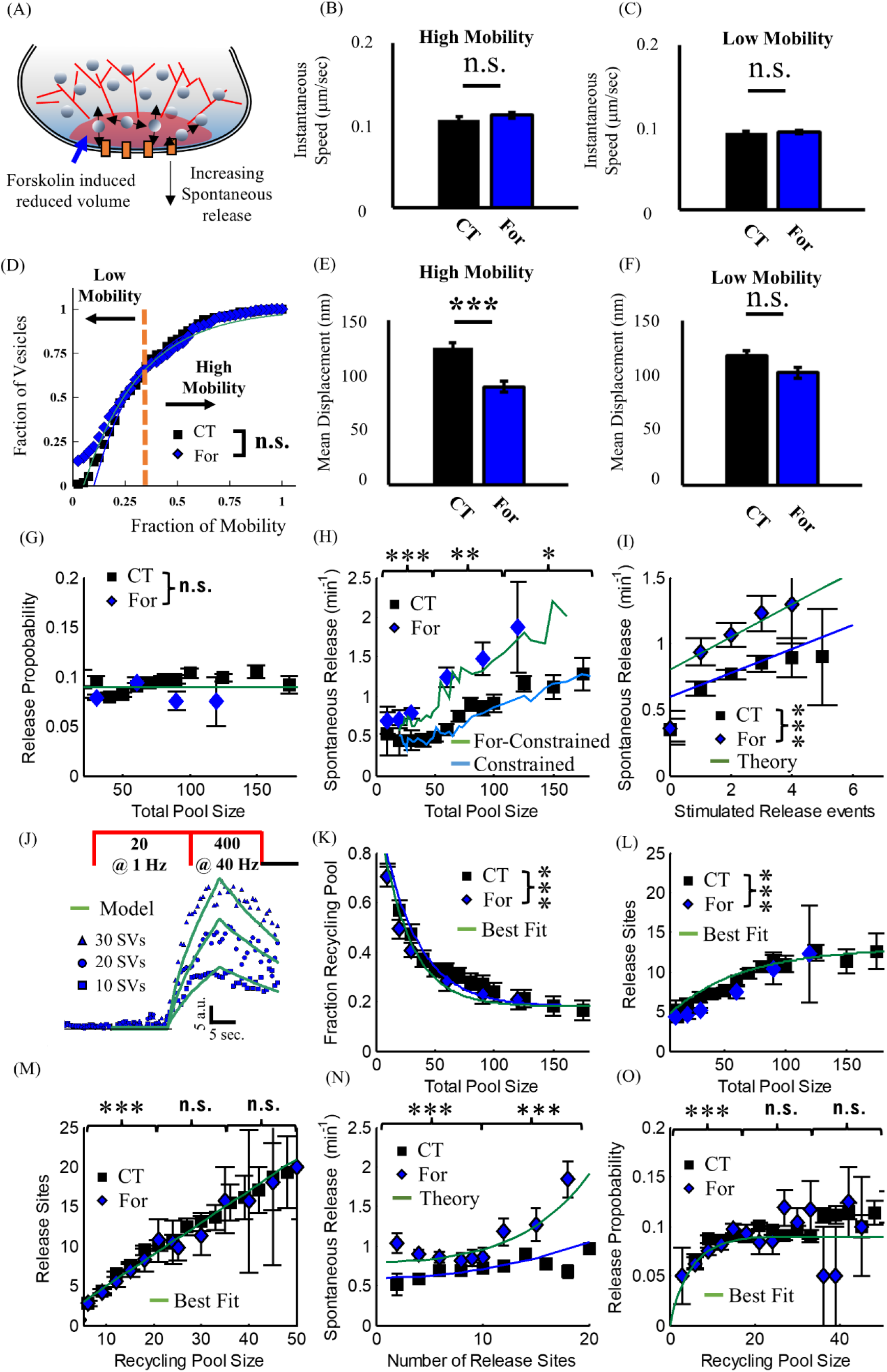
Forskolin Induced Reduction in Recycling Pool Volume Increases Spontaneous Release Rate (A) Cartoon representation of reduced Recycling Pool volume in the presence of Forskolin (For) leading to increasing spontaneous release rate. (B) Average High Mobility SV speed is compared for control (CT, Black) and in the presence of Forskolin (For, Blue). (C) Average Low Mobility SV speed is compared for control (CT, Black) and in the presence of Forskolin (For, Blue). (D) The distribution of SV mobility fraction in the presence of Forskolin (Blue Diamonds) has the same fraction of time mobile compared to control SVs (Black Squares). (E) Average High Mobility SV rms-displacement is significantly lower in the presence of Forskolin (For, Blue) compared for control (CT, Black). (F) Average Low Mobility SV rms-displacement is the same for control (CT, Black) and in the presence of Forskolin (For, Blue). (G) The average release probability is constant for all SV pool sizes (∼ 0.1) for both control (CT, Black Squares) and Forskolin (For, Blue Diamonds). (H) The spontaneous release rate increases more in the presence of Forskolin (Blue Diamonds), compared to control presynapses (Black Squares). The constrained computational model (Green Line) with decreased Recycling pool volume obtained from SGC5 measurements (Panel E) reproduces experimental For spontaneous release rates. (I) The spontaneous release rate increases linearly as a function of evoked release events for Forskolin (Blue Diamonds) presynapses, but with an overall higher scale factor compared to control (Black Squares). Both results follow the entropic force framework theory (Green and Blue lines), assuming a lower volume for the Recycling Pool in the presence of Forskolin. (J) Example traces of Forskolin presynapse pHluorin-VGlut1 intensity during 40Hz stimulation fit using the multinomial model (Green lines) for 10, 20, 30 SVs. (K) The fraction of SVs in the Recycling pool relative to the total number of SV pool is unchanged in the presence of Forskolin (Blue Diamonds) compared to control presynapses (Black Squares). (L) The number of release sites as a function of total pool size is unchanged in the presence of Forskolin (Blue Diamonds) compared to control (Black squares). Fit line (Green Line) is exponential recovery. (M) The number of release sites as a function of Recycling pool size is unchanged in the presence of Forskolin (Blue Diamonds) compared to control (Black squares). (N) The spontaneous release rate as a function of the number of release sites is higher in the presence of Forskolin (Blue Diamonds) compared to control presynapses (Black Squares). The entropic force theory (Green line) reproduces spontaneous release rates for number of release sites greater than 10. (O) The release probability as a function of Recycling Pool size is unchanged in the presence of Forskolin (Blue Diamonds) compared to control presynapses (Black Squares). Fit line (Green Line) is exponential recovery. CT: pHluorin-VGlut1: N_presynapses_ = 2057, from 6 different cultures; SGC5: N_SVs_ = 171, from 5 different cultures; N_simulations_ = 100 per SV number Forskolin: pHluorin-VGlut1: N_presynapses_ = 457, from 3 different cultures; SGC5: N_SVs_ = 275, from 6 different cultures; N_simulations_ = 100 per SV number Statistics: 2-sampled KS-test performed in (D). For all other panels, pair-wise 2-sampled t-Test.* = P< 0.05; ** = P < 0.01; *** = P< 0.001. Data points represent average values from binned raw data. Bins were set at fixed values to obtain a minimum of 30 data points per bin. Error bars represent SEM of binned data.

We first determined if neurons exposed to Forskolin exhibited significant changes in SV mobility, as previously observed.^41^ We exposed cell cultures to imaging media containing 50 μM Forskolin for 10- 20 min (See Methods). We then labeled single SVs and quantified SV mobility following the same protocol for control samples. We observed no significant change in SV speed for low or high mobility SVs (**Fig. 4 B,C**). We also observed no significant change in the fraction of SV mobility compared to controls (**Fig. 4 D**). However, we observed a significant reduction in SV displacement of high mobility SVs following Forskolin treatment (**Fig. 4 E**). This reduction in mobility of ∼28% is equivalent to the previously observed reduction in SV volume in slices.^41^ Importantly, we did not observe an equivalent reduction in the displacement of low mobility SVs (**Fig. 4 F**). These results support previous observations that Forskolin induces a constraint on the volume of the recycling pool.

Based on the single SV mobility results, the entropic force framework predicts an increase in spontaneous release rate under the same SV pool size distributions. To test this, we used the pHluorin- VGlut1 approach to measure spontaneous release and stimulated release rate after exposure to Forskolin. We observed a significant increase in spontaneous release rate for all measured SV pool sizes (**Blue diamonds, Fig. 4H**), but no significant change in release probability (**Blue diamonds, Fig. 4G**). We used the entropic force theory (**eqn. 1**) to model the spontaneous release rate using the experimentally measured 28% reduction in SV mobility (**Fig. 4 E**) and found that the theory predicted the experimentally measured pHluorin-VGlut1 rate (**Solid blue line, Fig. 4H**). Moreover, comparing the spontaneous release to the number of stimulated release events in the same synapse showed a significant increase in the overall spontaneous rate for Forskolin (**Blue diamonds, Fig. 4I**) compared to control conditions (**Black Squares, Fig. 4I**). These results suggest that spontaneous release rates occur independent of any changes in active zone mechanics.

To confirm that the observed changes in spontaneous release rate were exclusively due to the reduction in SV volume, we also compared spontaneous and stimulated release to the recycling pool using our multinomial fit approach to the 40 Hz intensity (**Fig. 4J**). We observed no significant change in the relative fraction of recycling pool SVs (**Fig. 4K**) or in the number of release sites relative to the total pool size (**Fig. 4L**). We also observed no change in the number of release sites (**Fig. 4M**) or release probability (**Fig. 4O**) relative to the recycling pool size.

However, we did observe a significant increase in the spontaneous release rate with increasing number of release sites compared to control (**Fig. 4N**). The increase in spontaneous release rate relative to release sites suggests that the recycling pool is increasingly constrained in the presence of Forskolin for the same number of release sites leading to an increase in spontaneous release per release site, consistent with the entropic force framework.

*These combined results show that the entropic force model can predict changes in presynaptic spontaneous release rate based on changes in the SV pool dynamics*.

## Conclusion

In the present study, we used an innovative combination of established experimental and computational tools to show that the entropic force framework can model and predict spontaneous release frequency. We introduced the entropic force framework using La-SEM analysis of cultured hippocampal neurons (**Fig. 1**). We then used a computational model to show how the experimentally observed distribution of SVs within the presynapse leads to a spontaneous release rate. We used a pH-sensitive fluorescent reporter (pHluorin-VGlut1) to experimentally measure the spontaneous release rate (**Fig. 2**). We reproduce the observed spontaneous release rate using our computational and theoretical model. We further show that the active zone mechanics play a supporting role in the spontaneous release rate. We used an alternative reporter (SGC5) to label and experimentally measure SV mobility within the presynapse (**Fig. 3**). We show that SV mobility supports the hypothesis that the Reserve Pool SVs constrain the Recycling pool SVs. We then combine a bootstrapping approach of SGC5 data with the entropic force framework to reproduce the pHluorin-VGlut1 spontaneous release. Lastly, we show that the spontaneous release rate can be increased by restricting the volume of the Recycling pool using the agent Forskolin (**Fig. 4**). These combined results support the entropic force framework that the Reserve SV pool constrains the volume of the Recycling pool to exert of force on the RRP regulating the spontaneous release rate.

## Discussion

### Why does the entropic force framework predict spontaneous release rate?

The spontaneous release rate can vary widely among presynapses and appears to have little correlation with evoked release.^15,36,37^ Further, there are a large number of biological parameters, from the number of release sites to the presynaptic actin polymerization rate, that could be regulated and lead to different spontaneous release rates.^15,16,28^ This complexity has limited the ability to coherently understand what, if anything, regulates spontaneous release. However, the spontaneous nature itself provides the framework to model release rate. Spontaneous processes in any system are a competing balance between (i) the energy available to the system and (ii) the maximization of the possible configurations of the system (entropy). Presynapses generate an SV energy density by constraining the mobility of the Recycling pool, which in turn generates an entropic force due to the fact that the mobility of the Recycling SVs works to drive them away from each other. This increase in energy density and entropic force is then directed toward the RRP resulting in a spontaneous exocytosis event. This framework puts the complex set of biological parameters into a single dominant pathway. Any change in the presynapse that changes the Recycling pool SV density will result in a corresponding change in the force on the RRP and thus spontaneous release rate.

### What determines the homeostatic spontaneous release rate?

The entropic force theory combined with the experimental results in this study suggest that a dynamic equilibrium is maintained within the presynapse to maintain a spontaneous release rate while minimizing the energy required to do so. Our results show that the density of the SVs in the recycling pool is more important than the number. Our results also show that the number of the SVs in the recycling pool is typically kept around 20 +/- 5. We suggest that this is an equilibrium based on multiple constraining factors: (i) SV kinetic energy is limited by their diffusive speed, which is itself a constrained value (∼0.15 μm/sec); (ii) the dominant energy cost in any presynapse is maintaining homeostatic SV neurotransmitter concentration,^61^ and thus the energy demand per presynapse increases with the number of total SVs; (iii) increasing the number of Recycling Pool SVs increases the generated force that the Reserve pool SVs must then constrain requiring more Reserve Pool SVs; (iv) the number of RRP (and corresponding release sites) correlates with the Recycling pool density in order to translate the entropic force into spontaneous release rate. These constraints, and probably many others, suggest that presynapses keep the size of the Recycling pool and Reserve pool regulated in order to maintain a fixed energy density that balances the energy costs of maintaining the pool with the force that the regulates then spontaneous release rate.

### How does entropic force fit with neurodegeneration?

Our hypothesis of the detailed balance required to maintain spontaneous release rate also provides a pathway to understand how molecular level changes lead to neurodegeneration. If the rate of spontaneous release is regulated to maintain the pre/post synaptic connection, then a reduction in that rate will result in a degradation of the synaptic connection. The entropic force provides a coherent framework to understand how different molecular level changes can lead to the same outcome (neurodegeneration). If a biological change (via mutation, external factors, etc.) leads to a reduction in SV density in the Recycling pool, then the spontaneous release rate will decrease and in turn lead to a reduction in the synaptic connection strength.

### How does entropic force fit within synaptic plasticity?

Our hypothesis that the Recycling pool density is the key factor that mediates spontaneous release rate provides a mechanism for how spontaneous release can be regulated during synaptic plasticity. For example, spontaneous release rate has been shown to dynamically change during different forms of synaptic plasticity.^30,31^ The entropic force framework suggests that the spontaneous release rate change can occur by dynamically regulating the number of SVs in the recycling pool in order to increase the density. Alternatively, as shown with the Forskolin results, the volume of the Recycling Pool can be reduced leading to an increase in the density of SVs. Either pathway would lead to an increase in spontaneous release rate, via an increase in the entropic force.

## Methods

### Primary Rat/Mice Protocols

Timed pregnant rats were obtained from a commercially available vendor (Sprague Dawley, Charles River), and were held for at least 48 hours after arrival and before dissection procedures. After 48 hours, the dam was euthanized using CO2 asphyxiation following NIH guidelines. Pup brains (at age E19) were then dissected, and the hippocampal portion of each lobe were removed. Pups were assumed to be an equal of mix of genders.

Mice were created by crossing B6.Cg-Tg(Camk2a-tTA)1Mmay/DboJ (Stock No. 007004) and FVB- *Fgf14^Tg(tetO-MAPT*P301L)^*^4510^*^Kha^* (Stock No. 015815) initially purchased from Jackson’s Laboratory and bred in-house. All offspring were genotyped before postnatal day 4 (PND 4), and only transgene-negative mice were used for cell cultures in the current study. All mice were provided food and water ad libitum, and housed under 12:12 hour light-dark cycles in humidity and temperature-controlled rooms. Mice used for dissection were assumed to be of mixed gender.

We have previously used both the rats and mice species in the present study for the same experimental approaches,^45,46,59^ in order to support cross-species validation. The Auburn University Animal Care and Use Committee approved all experiments (protocol numbers: 2020-3715, 2020-3657, and 2020-3742). All protocols complied with the guidelines published in the NIH Guide for the Care and Use of Laboratory Animals.

### Cell Culture

Cells were plated on previously prepared astrocytes on prepared glass coverslips in Neuronal Growth Media (84% Minimum Essential Medium (Thermo Fisher) with 9.6% Donor Equine Serum (Hyclone), 2% 1M Glucose in MEM (Thermo Fisher), 0.5% Penicillin/Streptomycin (Thermo Fisher), 1% N2 supplement (Thermo Fisher), 1% Sodium Pyruvate (v), 2% 1M HEPES, by volume). Plated cells were then kept in an incubator until imaging. At 24 hours afte dissection, neuronal growth media was replaced by neurobasal media (96% Neurobasal-A Medium (Thermo Fisher), 2.5% B-27 supplement (Thermo Fisher), 0.3% Glutemax-1 (Thermo Fisher)), 1.2% Penicillin/Streptomycin (Thermo Fisher), by volume). Additional neurobasal media was added every 7 days.

### Transfection Protocol

All cell cultures transfected with vGlut1-pHluorin followed the same protocol. Cultures were transfected at 3 DIV by replacing neurobasal medium with fresh 1 mL neurobasal medium containing pHluorin-VGlut1 viral titer. After 48 hours, the transfection medium was removed and replaced with 1ml of fresh Neurobasal media without virus. Every 7 days another 0.5 ml of neurobasal media was replaced with fresh neurobasal.

### Fluorescence Microscopy Experimental Approach

<u>Microscope:</u> Samples were imaged on a custom-built microscope with Ti2 base (Nikon), a 100x oil objective, Orca-Flash 4 CMOS camera with 6.5 x 6.5 μm pixel size (Hamamatsu), and illuminated with a Sola LED light source. Excitation and Emission light were filtered using commercial cubes (GFP, Nikon). All experiments were performed with samples at physiological temperature 37°C within a whole-microscope incubator (OKO Labs) at DIV14–19.

<u>pHluorin-VGlut1 experiments:</u> During experiments, cultures were perfused with bath solution (140 mM NaCl, 2.5 mM KCl, 2 mM CaCl2, 4 mM MgCl2, 10 mM HEPES, 2 mM Glucose, 50 mM DL-AP5, 10 mM CNQX, pH adjusted to pH 7.4), consistent with previous VGlut1-pHluorin experiments.^17,45,50,52,59^ Solutions were heated using a temperature controller attached to a multi-line solution heater (Warner Instruments). Ammonium Chloride solutions contained the above solution mixture plus 50 mM NH_4_Cl and pH-balanced to 7.4.

<u>Single SGC5 experiments:</u> Experiments were performed with samples at physiological temperature 37°C and DIV14–19. Cultures were initially stimulated with a single paired pulse to initiate a single exocytosis event followed by 4 min perfusion of media (140 mM NaCl, 2.5 mM KCl, 2 mM CaCl2, 4 mM MgCl2, 10 mM HEPES, 2 mM Glucose, 50 mM DL-AP5, 10 mM CNQX, pH adjusted to pH 7.25), consistent with our previously established protocols.^11,46^

<u>Pharmacological agents:</u> Samples exposed to Forskolin were pre-incubated for 5-10 min in imaging media containing 50 μM concentration. Samples exposed to Lat-A were pre-incubated for 30 min in imaging media containing 30 μM concentration. Media was then replaced with imaging media and either the pHluorin-VGlut1 protocol or the single SGC5 labeling protocol was then followed.

### Large Area Scanning Electron Microscopy

Cells are prepared for La-SEM following our previously established protocol.^39^ Cell cultures were washed with cacodylate buffer (0.15 M cacodylate buffer and 2 mM CaCl2). Then, cells were aldehyde fixed for 15 minutes at 37°C in modified Karnovsky’s fixative (2.5% glutaraldehyde, 2% paraformaldehyde, and cacodylate buffer) and stored in fixative at 4 °C overnight. The next day, samples were washed with cacodylate buffer. A light-protected heavy metal incubation was then performed in a solution of 1.5% potassium ferrocyanide, 1% osmium tetroxide, and cacodylate buffer for 1 hour, followed by washing. Thiocarbohydrazide-osmium liganding (OTO) was performed via light-protected thiocarbohydrazide (aq) incubation at 60°C for 20 minutes, followed by a serial wash and a light-protected incubation in 2% osmium tetroxide (aq) for 30 minutes. Samples were washed and incubated in 1% uranyl acetate (aq) overnight at 4°C with light protection. The following day, samples were washed, and contrast was enhanced with en bloc staining in 20 mM lead aspartate at 60 °C for 30 minutes. Samples were again washed and underwent a stepwise dehydration series in ice-cold ethanol for 10 minutes each (Ethanol/ddH20: 20%, 50%, 70%, 90%, 100%, and 100%). After dehydration, samples underwent serial resin infiltration at 2-hour intervals (Durcupan/ethanol: 25%, 50%, and 75%). Samples were then placed in 100% Durcupan resin overnight and then into fresh 100% Durcupan resin with the accelerator. Cell coverslips are then placed on an aluminum weigh boat and cured in a 60 °C oven for 48 hours.

Post resin curing, samples were sent to the Washington University Center for Cellular Imaging (WUCCI) where the coverslips were exposed with a razor blade and etched off with concentrated hydrofluoric acid. Small pieces of the resin containing the cells were then cut out and mounted onto blank resin stubs before 50 nm thick five serial sections were cut in the cell culture growing plane and placed onto a silicon wafer chips. These chips were then adhered to SEM pins with carbon adhesive tabs and large areas (∼ 300 x 300 µm) were then imaged at high resolution in a FE-SEM (Zeiss Merlin, Oberkochen, Germany) using the ATLAS (Fibics, Ottowa, Canada) scan engine to tile large regions of interest. High-resolution tiles were captured at 16,384 x 16,384 pixels at 5 nm/pixel with a 5 µs dwell time and line average of 2. The SEM was operated at 8 KeV and 900 pA using the solid-state backscatter detector. Tiles were aligned using ATLAS 5.

<u>La-SEM Analysis</u>: Raw SEM images were loaded into ImageJ and filtered using ImageJ’s Gaussian Blur filter with a rolling ball radius of 1. The images were scanned in a raster form, starting from the top left. Presynapses were identified as intact regions with a grouping of SVs (>10). The area of each presynapse was found by tracing the perimeter and quantifying the ellipticity and total area. The active zone was identified as the darkest region along the perimeter where there were SVs within 50 nm. A *radial axis* was defined from the center of the active zone and extending perpendicularly along the presynapse (Black arrow, **Fig. 1B**). Lines were drawn at 50 nm intervals along the radial axis (*angular line*) and perpendicular to the radial axis (Colored lines, **Fig. 1B**), and extending to the perimeter of the presynapse. The total number of SVs at each 50 nm radial point were recorded along with the length of the *angular line*. The density of SVs at each 50 nm radial point was calculated as the total number of SVs along each line, divided by the total length of each line (with voids larger than 100 nm subtracted from the total length of the line). The SVs defined as RRP are the number of SVs at 50 nm from the active zone. The Recycling pool SVs were defined as the SVs within the first 200 nm of the active zone.

### Derivation of Spontaneous Release Model

In order to determine if the spontaneous release of vesicles was mediated by the macroscopic pool dynamics, we developed a framework to relate presynapse structure, pool dynamics, and release frequency. In this framework we assume that the pool dynamics are driven by entropic force, generated by vesicle diffusion, which means that the pressure the vesicles exert on the walls of the presynapse can be described similarly as done with the ideal gas law. In this model, vesicles in the pool are also constantly exerting force on the vesicles tethered at the active zone as well as each other. This applied diffusive force will result in some tethered vesicles occasionally fusing with the plasma membrane leading to an exocytosis event. We propose that the resulting spontaneous release frequency of vesicles is then mediated by the pressure exerted by the synaptic vesicle pool as follows:

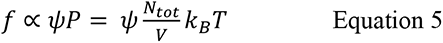

where N_tot_ is the total number of synaptic vesicles within the presynapse. The energy from each vesicle, which is what contributes to the force applied to the plasma membrane, is proportional to their speeds 𝑘_𝐵_𝑇 ∝ 𝑣^2^.

In this framework the **spontaneous correlation parameter** (y) represents the relationship between the spontaneous release frequency and: (i) the properties of the vesicles within the presynapse, (ii) the properties of the presynapse active zone release sites, and (iii) the local environment near the active zone (i.e. Ca^2+^ concentration).

First, we will establish how the properties of the vesicles can be separated into experimentally measurable parameters. It is well established that vesicles exist in different pools within the presynapse.^7,9^ The two major pools are: (i) the recycling pool, and (ii) the reserve pool. Further, vesicle mobilities (displacement) are different within the pools,^9^ with vesicle mobility in the reserve pool being lower due to being tethered to each other and to the actin network (Fig. 1).^48,62,63^ We then separate the relative contributions of each pool to the spontaneous release rate:

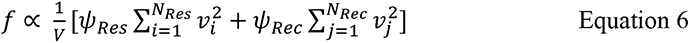

where both N_Res_ and N_Rec_ are both functions of N_Tot_ through conservation of vesicles (N_Tot_ = N_Rec_ + N_Res_, Fig. 1).

Second, we will now establish how the properties of the presynapse can be separated into experimentally measurable parameters for the **spontaneous correlation parameter.** The active zone and vesicle release sites play essential roles in determining the probability of vesicle exocytosis.^2,16^ This means that more release sites will result in higher spontaneous release frequency. Here we assume that the contribution of SV force on spontaneous release is also a function of the number of release sites:

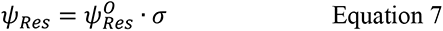

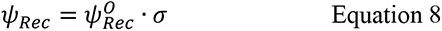

where σ is the relationship between the number of release sites and the number of vesicles in the presynapse. We use experimental results to show that this is dependent upon the total pool size. Thus, the full relationship between the spontaneous release frequency and the pressure from the vesicles within the presynapse is given by:

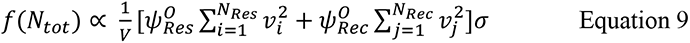

Finally, we explicitly establish the relationship between the volume itself and its effect on the spontaneous release frequency. In general, presynapse size scales with the number of vesicles across different types of synapses. In the present study, we chose to quantitatively distinguish this relationship in central synapses using our previously established La-SEM approach. We then counted the number of vesicles within the presynapse and quantified the relationship between presynapse volume and the number of vesicles (**Fig. 1C**). Presynapse volume increased linearly with the number of vesicles resulting in:

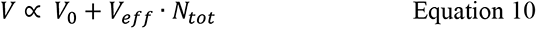

where V_0_ is the smallest possible presynapse volume and V_eff_ is the volume increase per vesicle in the presynapse.

Lastly, the Reserve pool SVs are tethered to each other and the actin network away from the active zone. This means that they do not significantly contribute to the entropic force applied to SVs at the active zone. In our framework they would thus not contribute to the spontaneous release rate. Consequently, the spontaneous release rate equation reduces to:

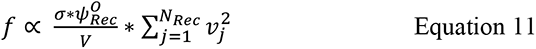

where the volume now represents the constrained volume of the Recycling pool.

### Work-flow of pHluorin-VGlut1 data analysis

All pHluorin-VGlut1 data was analyzed using the same approach using imageJ and custom written MATLAB code as follows:

i. Raw tiff files were background subtracted using a 10-pixel rolling-window algorithm in imageJ.
ii. Bright intensity puncta were identified at frame 1080 (at pulse 300 of 400 total) in imageJ
iii. Identified bright puncta locations were integrated with a 16x16 pixel area and the time-dependent integrate intensity was plotted
iv. Presynapses were selected for analysis if they exhibited intensity response at both 40 Hz stimulation and NH_4_Cl exposure (Fig. 2C), regardless of whether they exhibited spontaneous or evoked release.
v. The spontaneous release and stimulated release events and corresponding intensities were then identified as described below.
vi. The maximum NH_4_Cl intensity was then identified as the mean value of the top 10 intensity frames. The intensity was converted to number of vesicles by dividing by the mean single pHluorin-VGlut1 intensity per vesicle.
vii. The 40 Hz intensity was then fit using the multinomial release model described below, based on the mean intensity per single vesicle obtained in step (vi). Note, the mean single vesicle intensity was calculated on a sample-by-sample basis.
viii. All resulting values were then combined on a synapse-by-synapse basis and correlation plots were made as shown in each figure.

### Spontaneous and Stimulated Release Identification from pHluorin-VGlut1

Each presynaptic integrated intensity was run through a Kenedy-Chung filtering algorithm following established published parameters for VGlut1-pHluorin experiments.^50^ The resulting distribution of background intensities were then fit using a normal distribution to determine the first standard deviation of noise (σ in Fig. 2C). Single vesicle release events were then identified as any change in intensity greater than one standard deviation of the background that occurs within 500 msec or less (single events in Fig. 2C). The intensity for release event was calculated as the peak intensity minus the 3 frame average background intensity prior to the release event. The total time of intensity observed was calculated as the time from beginning of rising intensity to the time the intensity decreased below one standard deviation of the noise.

### Fitting 40 Hz pHluorin-VGlut1 Intensity

i. <u>Multinomial computational model</u>: We used our previously established computational model to fit our 40Hz data,^45^ which is based on the established multinomial model of presynaptic transmission.^36,37,64^ First, the model predicts a presynaptic response to a single stimulus as: R = n*p*q where R is the total response from the presynapse, p is the probability of release per vesicle to a single stimulus, n is the number of active zone sites, and q is the post-synaptic response to a single vesicle release. Here we modify the traditional model to reproduce pHluorin-VGlut1 intensity as follows:

a. R = I(t) = the total pHluorin-VGlut1 intensity as a function of time
b. q = i = the intensity per vesicle that undergoes exocytosis and then decays with time due to endocytosis
c. p is a probability threshold used for stochastic release comparison The pHluorin-VGlut1 intensity during a train of stimulation is then given by:

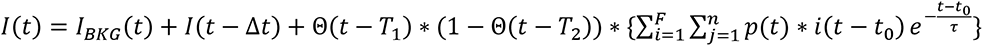

where each simulation time-step (Δt) includes the intensity from the previous simulation time-step; a total background intensity (I_BKG_); a probabilistic increase in intensity given by a per vesicle count (i) from each stimulus pulse; each vesicle has the same endocytosis rate (τ); F is the number of stimulation pulses per time-step; n is the number of release sites at the active zone; and Θ is the Heaviside function that is zero before the time (T) and unity after the time (T).
ii. The equation is computationally generated using the following algorithm:

a. An intensity array is generated for the entire time-window of the modeled data
b. The background intensity is calculated as a function of time and added to the array at each time-step
c. The stimulation time-window is designated, and the frequency of stimulation is predefined
d. During the stimulation time-window, a release event is stochastically calculated using a threshold
iii. <u>Generation of intensity profile database and fitting</u>: In order to fit each experimental 40 Hz intensity profile, a database of modeled profiles is created using the algorithm described above. The database was created by varying each parameter in the model over the range of values and steps listed in **table S1**. Each experimental 40 Hz intensity profile was then compared to every modeled profile in the database. The quality of fit for each profile was determined by a chi-square fitting algorithm. A single modeled profile was chosen with the lowest chi-square comparison to the experimental 40 Hz profile. The parameters from the modeled profile (number of release sites, total recycling pool, etc…) was then combined with the single and spontaneous release events for the specific presynapse resulting in a complete presynapse database.

### Computational Simulation Model of SV mobility

<u>Presynapse structure:</u> Simulated presynaptic vesicle pool mobility was performed using a lattice-based model approach in python. An **n**\***m** array is generated for each presynapse representing all spatial locations within the presynapse. Each lattice site is designated as 50x50 nm in area, equal to the size of a single SV. Each time-step in the simulation is designated as 300 msec. The active zone is designated at x=0 of the presynapse array. The number of n-lattice sites at x = 0 is then designated as equal to the number of release sites. The size of the presynapse is constrained as follows: (i) n = m for presynapses less than 10 lattice sites (50 nm); (ii) for all m > 10, n = fixed at 10 lattice sites (equal to a maximum of 10 release sites at the active zone).

<u>Vesicle Starting Distributions:</u> A starting distribution of reserve pool vesicles is generated as either (i) random for the unconstrained model, or (ii) at fixed locations starting at 5 lattice sites from the designated active zone. A starting distribution of recycling pool vesicles is then generated at random locations within the first 5 lattice sites from the designated active zone (x = 0). Vesicles are then randomly assigned starting velocities as either (i) zero for reserve pool vesicles in the constrained model; (ii) equal to one lattice site in x and/or y for recycling pool vesicles; (iii) equal to one lattice site in x and/or y for reserve pool vesicles in the unconstrained model. The number of vesicles in each pool is determined based on the experimentally determined relationship obtained using La-SEM and pHluorin-VGlut1 (**Fig. S1 A**). Each vesicle is assigned a number (1, 2, 3,…,N) up to the total number of vesicles in the simulation.

Vesicle Mobility Rules: During simulations, vesicle distributions within the presynaptic array are updated using the following rules: (i) The presynaptic lattice is updated with the current position of each vesicle; (ii) each SV position is updated in chronological order (1, 2, 3 …); (iii) the next position of the SV is calculated based on the location of the previous position plus the velocity of the SV; (iv) if the next position is occupied, then SV is assumed to engage in an elastic collision with the SV in the next lattice site; (v) the SV velocity is reversed and the next lattice site is calculated based on the previous lattice site plus the new reversed velocity; (vi) if the new position is also occupied, then the next neighboring lattice sites are all checked for occupancy; (viii) if all neighboring lattice sites are occupied, then the SV does not move for the current time-step and is re-checked on the next time-step; (ix) if the SV reaches the edge of the lattice (x = **m**, or y = 0 or **n**) the SV elastically collides with the membrane and reverses direction; (x)

<u>Spontaneous vesicle exocytosis rules:</u> if the next lattice site that an SV will occupy is the active zone (x = 0), then a random number is generated between (0,1); if the random number is below a fixed threshold (p = 0.002 per hit), then the SV engages in an exocytosis event which is recorded for that time-step. At the end of each simulation, the average time between recorded exocytosis events is recorded as a rate per min. After 100 simulations, the average spontaneous rate per minute is calculated along with SEM for the rate.

### Single SV Data Analysis

Single SV identification both the single and bulk raw tiff files were performed in imageJ and identified as bright puncta in the single movie that also had a larger corresponding bright puncta in the bulk movie (defined as a bulk intensity puncta >2 of the single movie puncta intensity). The position for each single SV and presynapse were recorded. Single SVs were secondarily confirmed using established computational track identification algorithms.^11,14,46,65^ High-resolution SV track positions were then run through our previously established correlation analysis algorithm to identify bouts of Fast, Diffusive, and Pausing behavior.^11,46^ The SV speed was determined as the average speed per frame averaged across all frames in a track. The average speed per condition is then calculated as the average speed across all tracks. The displacement was calculated as the average mean-square-displacement between 400 – 700 frames for each track calculated from the track position in the first frame. The displacement for each condition is then calculated as the average of all tracks measured for each condition.

### Statistical Analysis

KS-tests were used for cumulative distribution comparisons. For all other measurements, statistical analysis followed the same approach. First, all data were compared using pair- wise t-Test for each measurement and condition (i.e. all CT spontaneous release rates were compared to all Lat-A spontaneous release rates) to determine if there were any statistical significance. Second, if significance was observed in the whole data set, then data were compared for specific ranges (i.e. different SV pool size ranges in **Fig. 2F**) to determine which ranges exhibited significant differences. Third, all binned data points were compared on a bin-by-bin basis, for the same bins (i.e. the spontaneous release rates were compared on a bin-by-bin basis for each release site bin in **Fig. 2M**). Statistical results were reported in each measurement for total data and/or data ranges where significances were observed.

## Author Contributions

RC and MR maintained mouse animal colonies and performed mouse colony cell culture preparations. MP performed rat cell culture preparations. PW and MP performed single SGC5 experiments and MWG performed VGlut1-pHluorin and La-SEM experiments. PW analyzed the pHluorin-VGlut1 data and La- SEM data and was blind to conditions. MWG, analyzed the SGC5 data and was blind to conditions. MWG and NK developed computational models of synaptic vesicle mobility and spontaneous release rates. MWG and MR designed the study and obtained funding for the study. All authors contributed to the writing and editing of the manuscript.

## Conflicts of Interest

The authors report no conflicts of interest.

## Acknowledgements

The authors would like to acknowledge funding for this study from Auburn University internal grant program. The authors would also like to acknowledge the Washington University Viral Vector Core in the Hope Center for pHluorin-VGlut1 viral vector production. We acknowledge the Washington University Center for Cellular Imaging (WUCCI) for the La-SEM measurements. Finally, the authors would like to acknowledge that *no part* of this study design, implementation, code development, data analysis, manuscript writing, or figure generation involved the use of artificial intelligence, and all work is the authors own.

---

**Supplementary Figure 1:**
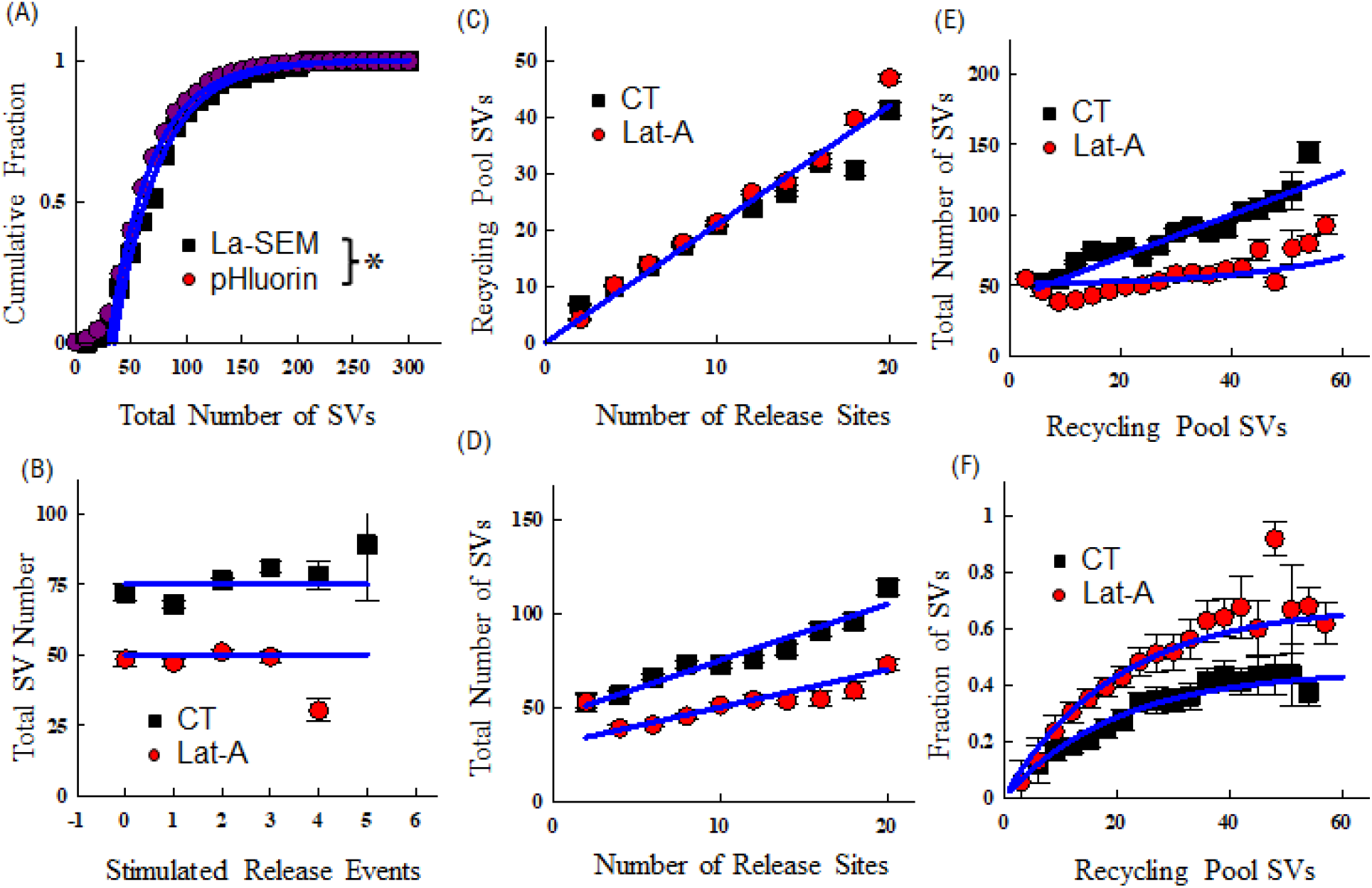
(A) Distribution of total number of SVs per synapse measured with pHluorin-VGlut1 and La-SEM. Data were fit to a cumulative fraction (Blue Lines). (B) Average total number of SVs measured as a function of stimulated release events on a synapse-by- synapse basis for CT (Black Squares) and Lat-A (Red Circles). Data were fit for an average value across all release event bins (Blue Lines). (C) Average number of Recycling pool SVs as a function of release sites on a synapse-by-synapse basis for CT (Black Squares) and Lat-A (Red Circles). Data were fit to a line (Blue lines). (D) Average number of total SVs measured as a function of the number of release sites on a synapse-by- synapse basis for CT (Black Squares) and Lat-A (Red Circles). Data were fit to a line (Blue Lines). (E) The average total number of SVs measured as a function of number of recycling pool SVs on a synapse-by-synapse basis for CT (Black Squares) and Lat-A (Red Circles). (F) The fraction of SVs in the recycling pool as measured as a function of number recycling pool SVs on a synapse-by-synapse basis for CT (Black Squares) and Lat-A (Red Circles).

**Table Supplementary 1:** Multinomial Fits Parameters

| Parameter | Lower Limit | Upper Limit | Step-Size | Units |
| --- | --- | --- | --- | --- |
| n_sites | 1 | 22 | 1 | N/A |
| Response_start | 0 | 9.9 | 0.5 | seconds |
| Response_end | 0.1 | 10 | 0.5 | seconds |
| tau_endocytosis | 15 | 25 | 0.25 |  |
| Intensity | 350 | 600 | Set by experiment | cnts/sec |
| Fixed Parameters |  |  |  |  |
| P_site | 0.01 | 0.01 | 0 | Probability |
| P_MVR | 2 | 2 | 0 | Fraction |
| P_MVR_decay | 2 | 2 | 0 | seconds |
| Stimulations Per second | 40 | 40 | 0 | N/A |
| Total SV Pool Size | 100 | 100 | 0 | N/A |

## Notes

### Competing Interest Statement

The authors have declared no competing interest.

## References

1. Rizo, J. & Rosenmund, C. Synaptic vesicle fusion. Nat. Struct. Mol. Biol. 15, 665–674 (2008).

2. Gramlich, M. W. & Klyachko, V. A. Nanoscale Organization of Vesicle Release at Central Synapses. Trends Neurosci. 42, 425–437 (2019).

3. Tucker, W. C., Weber, T. & Chapman, E. R. Reconstitution of Ca2+-Regulated Membrane Fusion by Synaptotagmin and SNAREs. Science 304, 435–438 (2004).

4. Maschi, D. & Klyachko, V. A. Spatiotemporal Regulation of Synaptic Vesicle Fusion Sites in Central Synapses. Neuron 94, 65–73.e3 (2017).

5. Südhof, T. C. The Presynaptic Active Zone. Neuron 75, 11–25 (2012).

6. Denker, A., Kröhnert, K., Bückers, J., Neher, E. & Rizzoli, S. O. The reserve pool of synaptic vesicles acts as a buffer for proteins involved in synaptic vesicle recycling. Proc. Natl. Acad. Sci. 108, 17183 (2011).

7. Alabi, A. A. & Tsien, R. W. Synaptic vesicle pools and dynamics. Cold Spring Harb. Perspect. Biol. 4, a013680–a013680 (2012).

8. Chanaday, N. L., Cousin, M. A., Milosevic, I., Watanabe, S. & Morgan, J. R. The Synaptic Vesicle Cycle Revisited: New Insights into the Modes and Mechanisms. J. Neurosci. 39, 8209 (2019).

9. Rizzoli, S. O. Synaptic vesicle recycling: steps and principles. EMBO J. 33, 788–822 (2014).

10. Rizzoli, S. O. & Betz, W. J. Synaptic vesicle pools. Nat. Rev. Neurosci. 6, 57–69 (2005).

11. Forte, L. A., Gramlich, M. W. & Klyachko, V. A. Activity-Dependence of Synaptic Vesicle Dynamics. J. Neurosci. 37, 10597 (2017).

12. Kamin, D. et al. High- and Low-Mobility Stages in the Synaptic Vesicle Cycle. Biophys. J. 99, 675–684 (2010).

13. Denker, A. & Rizzoli, S. Synaptic Vesicle Pools: An Update. Front. Synaptic Neurosci. 2, 135 (2010).

14. Park, H., Li, Y. & Tsien, R. W. Influence of Synaptic Vesicle Position on Release Probability and Exocytotic Fusion Mode. Science 335, 1362–1366 (2012).

15. Farsi, Z., Walde, M., Klementowicz, A. E., Paraskevopoulou, F. & Woehler, A. Single synapse glutamate imaging reveals multiple levels of release mode regulation in mammalian synapses. iScience 24, 101909 (2021).

16. Ralowicz, A. J., Hokeness, S. & Hoppa, M. B. Frequency of Spontaneous Neurotransmission at Individual Boutons Corresponds to the Size of the Readily Releasable Pool of Vesicles. J. Neurosci. 44, e1253232024 (2024).

17. Leitz, J. & Kavalali, E. T. Fast retrieval and autonomous regulation of single spontaneously recycling synaptic vesicles. eLife 3, e03658 (2014).

18. Fernández-Alfonso, T. & Ryan, T. A. The Kinetics of Synaptic Vesicle Pool Depletion at CNS Synaptic Terminals. Neuron 41, 943–953 (2004).

19. Maschi, D. & Klyachko, V. A. Spatiotemporal Regulation of Synaptic Vesicle Fusion Sites in Central Synapses. Neuron 94, 65–73.e3 (2017).

20. Gundlfinger, A., Breustedt, J., Sullivan, D. & Schmitz, D. Natural Spike Trains Trigger Short- and Long-Lasting Dynamics at Hippocampal Mossy Fiber Synapses in Rodents. PLOS ONE 5, e9961 (2010).

21. Doussau, F. et al. Frequency-dependent mobilization of heterogeneous pools of synaptic vesicles shapes presynaptic plasticity. eLife 6, e28935 (2017).

22. Papaleonidopoulos, V., Trompoukis, G., Koutsoumpa, A. & Papatheodoropoulos, C. A gradient of frequency-dependent synaptic properties along the longitudinal hippocampal axis. BMC Neurosci. 18, 79 (2017).

23. Dobrunz, L. E. & Stevens, C. F. Heterogeneity of Release Probability, Facilitation, and Depletion at Central Synapses. Neuron 18, 995–1008 (1997).

24. Pozzo-Miller, L. D. et al. Impairments in High-Frequency Transmission, Synaptic Vesicle Docking, and Synaptic Protein Distribution in the Hippocampus of BDNF Knockout Mice. J. Neurosci. 19, 4972 (1999).

25. Byczkowicz, N., Ritzau-Jost, A., Delvendahl, I. & Hallermann, S. How to maintain active zone integrity during high-frequency transmission. Presynaptic Act. Zone Mol. Plast. Dis. 127, 61–69 (2018).

26. Tseng, H. & Martinez, D. The Frequency Preference of Neurons and Synapses in a Recurrent Oscillatory Network. J. Neurosci. 34, 12933 (2014).

27. Andreae, L. C. & Burrone, J. The role of spontaneous neurotransmission in synapse and circuit development. J. Neurosci. Res. 96, 354–359 (2018).

28. Peng, A., Rotman, Z., Deng, P.-Y. & Klyachko, V. A. Differential Motion Dynamics of Synaptic Vesicles Undergoing Spontaneous and Activity-Evoked Endocytosis. Neuron 73, 1108–1115 (2012).

29. Gaffield, M. A., Rizzoli, S. O. & Betz, W. J. Mobility of Synaptic Vesicles in Different Pools in Resting and Stimulated Frog Motor Nerve Terminals. Neuron 51, 317–325 (2006).

30. Weichard, I., et al. Fully-primed slowly-recovering vesicles mediate presynaptic LTP at neocortical neurons. Proc. Natl. Acad. Sci. 120, e2305460120 (2023).

31. Fitzjohn, S. M. et al. An electrophysiological characterisation of long-term potentiation in cultured dissociated hippocampal neurones. Neuropharmacology 41, 693–699 (2001).

32. Ninan, I., Liu, S., Rabinowitz, D. & Arancio, O. Early presynaptic changes during plasticity in cultured hippocampal neurons. EMBO J. 25, 4361–4371 (2006).

33. Wasser, C. R. & Kavalali, E. T. Leaky synapses: Regulation of spontaneous neurotransmission in central synapses. Neuroscience 158, 177–188 (2009).

34. Andreae, L. C. & Burrone, J. The role of spontaneous neurotransmission in synapse and circuit development. J. Neurosci. Res. 96, 354–359 (2018).

35. Chen, S., et al. Real-time three-dimensional tracking of single vesicles reveals abnormal motion and pools of synaptic vesicles in neurons of Huntington’s disease mice. iScience 24, (2021).

36. Scheuss, V. & Neher, E. Estimating Synaptic Parameters from Mean, Variance, and Covariance in Trains of Synaptic Responses. Biophys. J. 81, 1970–1989 (2001).

37. Lanore, F. & Silver, R. A. Extracting quantal properties of transmission at central synapses. Neuromethods 113, 193–211 (2016).

38. Wang, B. & Dudko, O. K. A theory of synaptic transmission. eLife 10, e73585 (2021).

39. Maschi, D., Gramlich, M. W. & Klyachko, V. A. Myosin V functions as a vesicle tether at the plasma membrane to control neurotransmitter release in central synapses. eLife 7, e39440 (2018).

40. Lamanna, J., Gloria, G., Villa, A. & Malgaroli, A. Anomalous diffusion of synaptic vesicles and its influences on spontaneous and evoked neurotransmission. J. Physiol. 602, 2873–2898 (2024).

41. Rey, S., Marra, V., Smith, C. & Staras, K. Nanoscale Remodeling of Functional Synaptic Vesicle Pools in Hebbian Plasticity. Cell Rep. 30, 2006–2017.e3 (2020).

42. Masahiro Kaneko & Tomoyuki Takahashi. Presynaptic Mechanism Underlying cAMP-Dependent Synaptic Potentiation. J. Neurosci. 24, 5202 (2004).

43. Otmakhov, N. et al. Forskolin-Induced LTP in the CA1 Hippocampal Region Is NMDA Receptor Dependent. J. Neurophysiol. 91, 1955–1962 (2004).

44. Gobert, D. et al. Forskolin induction of late-LTP and up-regulation of 5′ TOP mRNAs translation via mTOR, ERK, and PI3K in hippocampal pyramidal cells. J. Neurochem. 106, 1160–1174 (2008).

45. Taipala, E., Pfitzer, J. C., Hellums, M., Reed, M. N. & Gramlich, M. W. rTg(TauP301L)4510 mice exhibit increased VGlut1 in hippocampal presynaptic glutamatergic vesicles and increased extracellular glutamate release. Front. Synaptic Neurosci. 14, (2022).

46. Parkes, M., Landers, N. L. & Gramlich, M. W. Recently recycled synaptic vesicles use multi- cytoskeletal transport and differential presynaptic capture probability to establish a retrograde net flux during ISVE in central neurons. Front. Cell Dev. Biol. 11, (2023).

47. Gramlich, M. W. & Klyachko, V. A. Actin/Myosin-V- and Activity-Dependent Inter-synaptic Vesicle Exchange in Central Neurons. Cell Rep. 18, 2096–2104 (2017).

48. Bingham, D. et al. Presynapses contain distinct actin nanostructures. J. Cell Biol. 222, e202208110 (2023).

49. Takamori, S. et al. Molecular Anatomy of a Trafficking Organelle. Cell 127, 831–846 (2006).

50. Chanaday, N. L. & Kavalali, E. T. Optical detection of three modes of endocytosis at hippocampal synapses. eLife 7, e36097 (2018).

51. Pfitzer, J., et al. Troriluzole Rescues Glutamatergic Deficits, Amyloid and Tau Pathology, and Synaptic and Memory Impairments in 3xTg-AD Mice. J. Neurochem. In Press, (2024).

52. Voglmaier, S. M. et al. Distinct Endocytic Pathways Control the Rate and Extent of Synaptic Vesicle Protein Recycling. Neuron 51, 71–84 (2006).

53. Jayaprakash, B. & Ryan, T. Single-vesicle imaging reveals that synaptic vesicle exocytosis and endocytosis are coupled by a single stochastic mode. Proc. Natl. Acad. Sci. U. S. A. 104, 20576–81 (2008).

54. Coué, M., Brenner, S. L., Spector, I. & Korn, E. D. Inhibition of actin polymerization by latrunculin A. FEBS Lett. 213, 316–318 (1987).

55. Morales, M., Colicos, M. A. & Goda, Y. Actin-Dependent Regulation of Neurotransmitter Release at Central Synapses. Neuron 27, 539–550 (2000).

56. Sankaranarayanan, S., Atluri, P. P. & Ryan, T. A. Actin has a molecular scaffolding, not propulsive, role in presynaptic function. Nat. Neurosci. 6, 127–135 (2003).

57. Bleckert, A., Photowala, H. & Alford, S. Dual pools of actin at presynaptic terminals. J. Neurophysiol. 107, 3479–3492 (2012).

58. Malagon, G., Miki, T., Llano, I., Neher, E. & Marty, A. Counting Vesicular Release Events Reveals Binomial Release Statistics at Single Glutamatergic Synapses. J. Neurosci. 36, 4010 (2016).

59. Maschi, D., Gramlich, M. W. & Klyachko, V. A. Myosin V Regulates Spatial Localization of Different Forms of Neurotransmitter Release in Central Synapses. Front. Synaptic Neurosci. 13, (2021).

60. Wu, Y., Yeh, F. L., Mao, F. & Chapman, E. R. Biophysical characterization of styryl dye-membrane interactions. Biophys. J. 97, 101–109 (2009).

61. Pulido, C. & Ryan, T. A. Synaptic vesicle pools are a major hidden resting metabolic burden of nerve terminals. Sci. Adv. 7, eabi9027.

62. Bloom, O. et al. Colocalization of synapsin and actin during synaptic vesicle recycling. J. Cell Biol. 161, 737–747 (2003).

63. Lee, S. J. et al. Role of Actin Filament on Synaptic Vesicle Pooling in Cultured Hippocampal Neuron. Appl. Microsc. 48, 55–61 (2018).

64. Reid, C. A. & Clements, J. D. Postsynaptic expression of long-term potentiation in the rat dentate gyrus demonstrated by variance-mean analysis. J. Physiol. 518, 121–130 (1999).

65. Jaqaman, K. et al. Robust single-particle tracking in live-cell time-lapse sequences. Nat. Methods 5, 695–702 (2008).

